# Fgf9-Nolz-1-Wnt2 Signaling Axis Regulates Morphogenesis of the Lung

**DOI:** 10.1101/2022.08.10.503529

**Authors:** Shih-Yun Chen, Fu-Chin Liu

**Author notes:** Corresponding author: Fu-Chin Liu, Ph.D. Institute of Neuroscience National Yang Ming Chiao Tung University 155 Sec. 2, Li-Nong Street Taipei 11221 Taiwan 886-2-2826-7216.

## Abstract

Morphological development of the lung requires complex signal crosstalk between the mesenchymal and epithelial progenitors. Elucidating the genetic cascades underlying the signal crosstalk is essential to understanding the morphogenesis of the lung. Here, we have identified Nolz-1/Znf503 as a mesenchymal lineage-specific transcriptional regulator that plays a key role in lung morphogenesis. The null mutation of Nolz-1 resulted in a severe hypoplasia phenotype, including decreased proliferation of mesenchymal cells, aberrant differentiation of epithelial cells, and defective growth of epithelial branches. The deletion of Nolz-1 also downregulated the expressions of *Wnt2*, *Lef1*, *Fgf10*, *Gli3* and *Bmp4*. Mechanistically, we found that Nolz-1 regulated lung morphogenesis primarily through Wnt2 signaling. Loss of function and overexpression studies demonstrated that Nolz-1 transcriptionally activated *Wnt2* and downstream β-catenin signaling to control mesenchymal cell proliferation and epithelial branching. The Nolz-1-Wnt2 axis was also supported by evidence that exogenous Wnt2 could causally rescue defective proliferation and epithelial branching in *Nolz-1* knockout lungs. Finally, we have identified Fgf9 as an upstream regulator of *Nolz-1*. Collectively, Fgf9-Nolz-1-Wnt2 signaling represents a novel signaling axis in the control of lung morphogenesis. These findings are also relevant to lung tumorigenesis in which a pathological function of Nolz-1 is involved.

## Introduction

The morphogenesis of the lung is coordinated by complex genetic cascades that occur in endoderm-derived epithelia and mesoderm-derived mesenchyme (Morrisey and Hogan, 2010; Warburton et al., 2000; Zepp and Morrisey, 2019). Transcription factors that are selectively expressed in the mesenchyme or epithelium are important for lung development (Costa et al., 2001; Maeda et al., 2007). For example, Foxf1, Tbx4/5, and Hox5 are expressed specifically in mesenchymal cell lineages (Chapman et al., 1996; Hrycaj et al., 2015; Ustiyan et al., 2018). Deletion of the mesenchyme-specific genes *Foxf1*, *Tbx4/5,* or *Hox5* results in a prominent phenotype of reduced lung buds (Cardoso and Lu, 2006; Morrisey and Hogan, 2010). Foxf1, Tbx4/5, and Hox5 regulate not only tracheal mesenchyme differentiation, but also epithelial cells. Knockout mouse studies show that Foxf1, Tbx4/5, and Hox5 regulate lung development through Wnt2/2b and Bmp4 signaling (Arora et al., 2012; Hrycaj et al., 2015; Ustiyan et al., 2018).

Epithelium-specific transcription factors include Nkx2.1/TTF1, Foxa1/a2, Sox2 (Maeda et al., 2007). Deletion or overexpression of these epithelium-specific transcription factors results in deficits in branching morphogenesis, differentiation of epithelial cells and mesenchymal cells, along with altered BMP or Shh signaling molecules (Gontan et al., 2008; Minoo et al., 1999; Wan et al., 2005). Because deletion of mesenchyme-specific transcription factors could non-cell autonomously affect epithelial development and vice versa, it highlights the importance of signal crosstalk between the mesenchymal and epithelial compartments. The signal molecules Fgf, Wnt, Shh, and BMP have been shown to underlie mesenchyme-epithelium interactions in lung development (Han et al., 2016).

*Nolz-1* (also known as *Znf503* and *Zfp503*) is a murine member of the *nocA/elB/tlp-1* (NET) gene family. The NET family is an evolutionally conserved gene family that includes *Tlp-1* (*C. elegans*), *elbow* (elb)/no ocelli (noc) (*Drosophila*), Nlz1 and Nlz2 (*zebrafish*), *Nolz-1* (chick), and *Zfp703/Nolz-2* and *Zfp503/Nolz-1* (mouse) (Chang et al., 2004; Chen et al., 2020; Dorfman et al., 2002; Ji et al., 2009; Runko and Sagerstrom, 2003; Zhao et al., 2002). Members of the NET family contain three core motifs, including the button-head box, Sp, and C2H2 zinc-finger motifs (Nakamura et al., 2004; Pereira et al., 2016). Nolz-1 and its homologues have been shown to function as a transcriptional repressor by interacting with the Groucho corepressor. Elb is associated with Groucho via the FKPY motif, whereas the interacting domain of Nlz1/Nlz2 with Groucho is mapped to the region between the bottonhead box and the C2H2 zinc-finger motif (Pereira-Castro et al., 2013; Runko and Sagerstrom, 2004). Chick Nolz-1 has also been shown to interact with Grg5. Functionally, *Nolz-1* and its homologues play a significant role in the developmental regulation of patterning, cell fate specification, and differentiation of neuronal and non-neuronal systems. Tlp-1 determines the asymmetric cell of tail cells that responds to Wnt signals in *C. elegans* (Zhao et al., 2002). Elb/noc regulates the development of eyes, legs, and trachea in *Drosophila* (Dorfman et al., 2002). Nlz1 and Nlz2 are involved in the patterning of rhombomeres in the development of the hindbrain and the closure of the optic fissure in *zebrafish* (Brown et al., 2009; Runko and Sagerstrom, 2003). Chick Nolz-1 is required for the specification of motor neurons (Ji et al., 2009). Murine Nolz-1 regulates the formation of striatal subdivisions in the mouse forebrain (Chen et al., 2020). The functional diversity of the members of the NET family implicates important roles of NET proteins in the regulation of the developmental regulation of different cell types in organogenesis.

The *Nolz-1 Drosophila* orthologue *elb/noc* has been shown to regulate the development of the trachea, a respiratory organ in *Drosophila* (Dorfman et al., 2002). Given the importance of *elb/noc* in the specification of tracheal branches in *Drosophila*, Nolz-1, as an evolutionarily conserved NET protein, could play a role in mammalian lung development. Here, we first identified Nolz-1 as a mesenchyme-specific transcriptional regulator in mouse lung development. We then investigated the biological function of Nolz-1 in the regulation of lung morphogenesis.

## Results

### Nolz-1 expression was developmentally regulated in the mouse lung

To investigate the temporal expression profile of Nolz-1 protein in the developing mouse lung, we performed the time course study of Nolz-1 protein expression by Western blotting. The results showed that Nolz-1 protein was highly expressed at E12.5 and E14.5 and was down-regulated at E16.5. The expression of Nolz-1 was drastically decreased after birth and was barely detectable in adult mouse lungs (Figure 1A). These findings suggest that Nolz-1 may play a role in the early stages of lung development.

**Figure 1.**
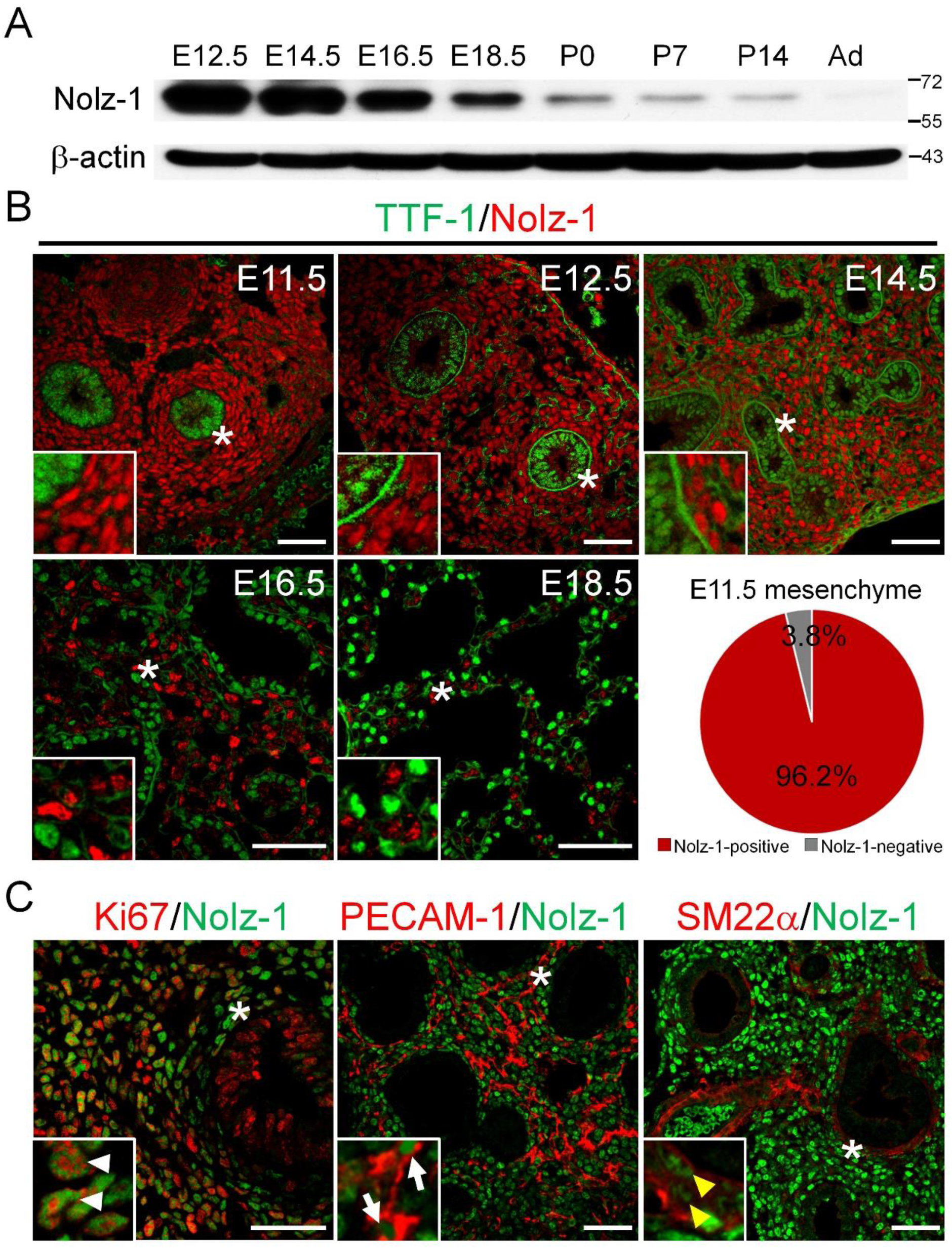
Spatiotemporal expression pattern and cell type-specific expression of Nolz-1 in developing mouse lungs. (A) Western blotting shows that Nolz-1 is developmentally regulated in the developing lung, with high levels at E12.5-E16.5 and gradually down-regulated after E16.5. Nolz-1 is barely detectable in the adult lung. (B) Nolz-1 is expressed specifically in the compartment of mesenchymal cells. Nolz-1 is expressed in mesenchymal cells in which the TTF1-positive epithelium is embedded. Nolz-1 (red) is not expressed in the TTF1-positive epithelium at E11.5, E12.5, E14.5, E16.5 and E18.5. The insets in b-f show high magnification of the regions indicated by asterisks. Nolz-1 is expressed in 96% of TTF1-negative mesenchymal cells at E11.5. (C) Nolz-1 is expressed in Ki67-positive proliferating cells in the E12.5 mesenchyme (arrowhead). Nolz-1 is co-localized with the endothelial cell marker PECAM-1 (arrow) and the smooth muscle marker SM22α (yellow arrowhead) in the E14.5 mesenchyme. E: embryonic day; P: postnatal day; Ad: adult. Scale bar in B, C, 50 μm.

### Nolz-1 was specifically expressed in the mesenchyme of embryonic lungs

To investigate the expression pattern of Nolz-1 protein, we performed immunostaining of Nolz-1. Consistent with the Western blotting results, we found that Nolz-1 protein was detected in the pseudoglandular (E12.5-E16.5), canalicular and saccular (E17.5-E18.5) stages of lung development (Costa et al., 2001; Maeda et al., 2007; Morrisey and Hogan, 2010; Warburton et al., 2000). Nolz-1 immunoreactivity was localized in the nucleus (Figure S1). Double immunostaining of Nolz-1 and Thyroid transcription factor 1 (TTF1/Nkx2.1), a marker of epithelial cells (Bohinski et al., 1994), showed that Nolz-1 was not co-localized in TTF1-positive epithelial cells at E11.5, E12.5, E14.5, E16.5 and E18.5 (Figure 1B). Nolz-1 was expressed in the majority of TTF1-negative mesenchymal cells at E11.5 (96 ± 0.7%; Figure 1B). These findings indicated that Nolz-1 was specifically expressed in the TTF1-negative mesenchyme of developing lungs.

### Nolz-1 was expressed in proliferating mesenchymal progenitor cells

We next determine whether Nolz-1 was expressed in proliferating progenitors of the mesenchyme. Double immunostaining of Nolz-1 and Ki67, a proliferating marker, showed colocalization of Nolz-1 and Ki67 by ∼70% in mesenchymal cells at E12.5 (Figure 1C). Nolz-1 was co-expressed with PECAM-1, a marker of progenitors and differentiated endothelial cells (White et al., 2007), in E12.5 mesenchymal cells (Figure 1C). Nolz-1 was co-expressed with SM22α, a marker of progenitors and differentiated smooth muscle cells (Goss et al., 2011; Solway et al., 1995), in E12.5 mesenchymal cells (Figure 1C). These findings identified Nolz-1 as a novel marker for mesenchymal cells in the developing mouse lung.

### Pulmonary hypoplasia in Nolz-1 null mutant embryos

To investigate the function of Nolz-1 in lung development, we studied *Nolz-1* germline knockout (KO) mice (Figure S1). *Nolz-1* KO mice died immediately after birth, probably due to lung failure, because lung size was dramatically decreased in E18.5 KO mice. Morphologically, the mutant lung had five lobes with proper orientations, but the mutant lobes collapsed and were significantly smaller than those of the wild type (Figure 2A). The time-course study showed that mutant lungs were morphologically normal at E11.5 compared to the wild type lungs. The impaired growth of mutant lungs became evident at E12.5, as the number of epithelial branches was reduced in mutant lungs (Figure 2B). By E14.5 and E16.5, the size of mutant lungs was about one-third of that of wild type lungs. The mutant lungs were much more compact and smaller than wild type lungs at E18.5.

**Figure 2.**
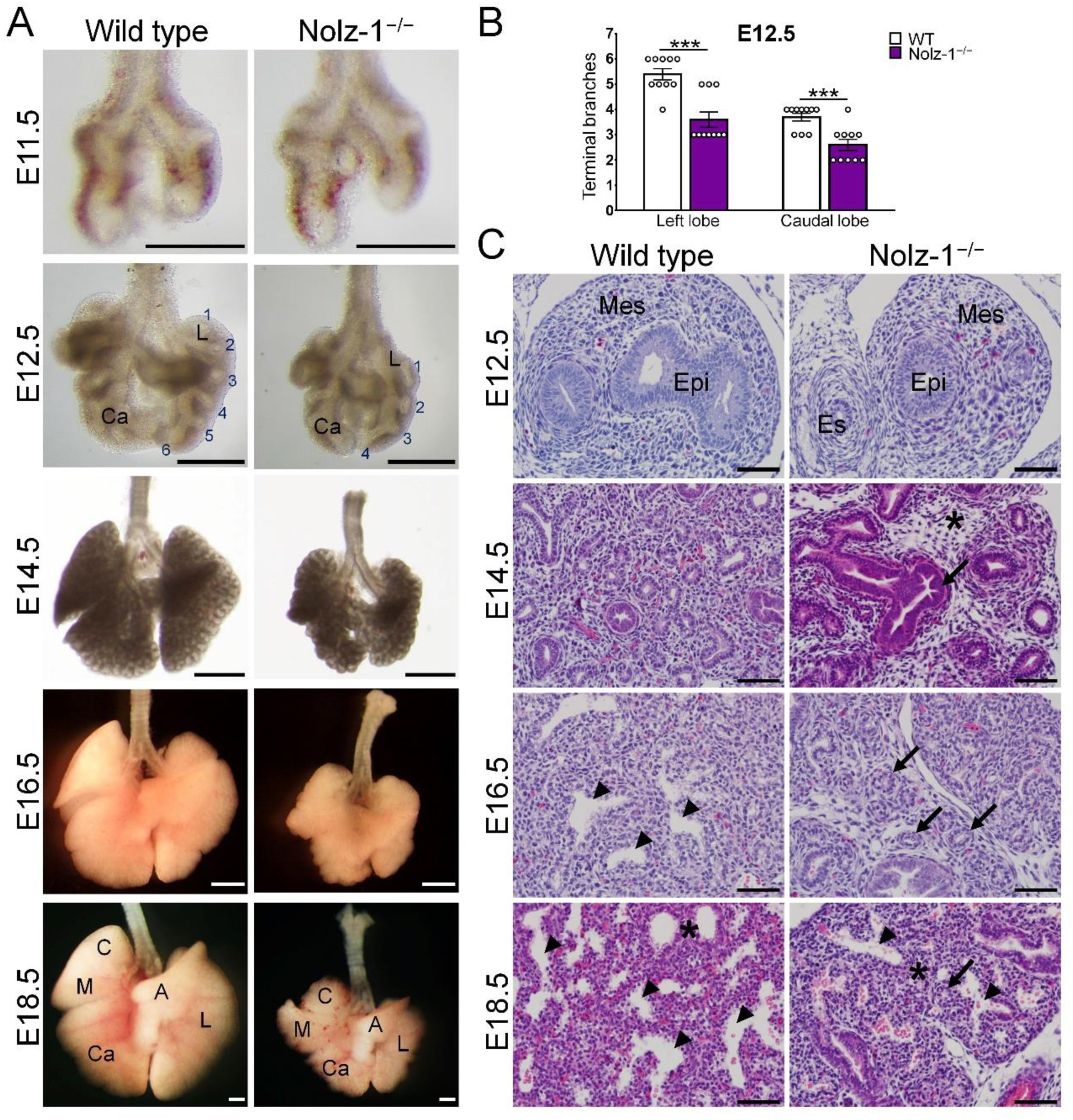
Hypoplasia of *Nolz-1* null mutant lungs. (A) The null mutation of *Nolz-1* does not affect the patterning of the lung buds into five lobules of the lung, but the overall size of the mutant lungs is significantly reduced from E12.5, E14.5, E16.5 to E18.5. (B) The numbers of epithelial branches are decreased in the left and caudal lobes of the mutant lungs. Student’s *t*-test, \*\*\**P* < 0.001, n = 10. (C) Hematoxylin and eosin staining. Aberrantly enlarged epithelial structures (arrow) and decreased mesenchymal cells (asterisk) are detected in E14.5 mutant lungs. Air sacs (arrowheads) are formed in wild type lungs at E16.5-E18.5, but few air sacs are present in mutant lungs. The epithelial columns (arrows) remained in mutant lungs. The air sac septation (asterisks) is increased in E18.5 mutant lungs. A: accessory lobe; C: cranial lobe Ca: caudal lobe; Epi: epithelium; Es: esophagus; L: left lobe; M: middle lobe; Mes: mesenchyme. Scale bars in A, 500 μm; C, 50 μm.

### Aberrant cytoarchitecture in developing *Nolz-1* null mutant lungs

Hematoxylin and eosin (H&E) stains showed that the mesenchyme and epithelium appeared to be similar between wild-type and KO lungs at E12.5. By E14.5, reduced and less compacted mesenchyme was observed in mutant lungs. Notably, enlarged mutant epithelial tubules were found at E14.5, but this phenotype was not observed after E14.5. At E16.5, which was the beginning of the canalicular stage, the epithelium began to form the saccular structure in wild-type lungs. By contrast, immature epithelial columns remained in mutant lungs at this stage. Furthermore, the mesenchymal septation between the air sacs was thicker in the mutant lungs than in the wild-type lungs (Figure 2C).

### Absence of abnormal apoptosis at early stages of *Nolz-1* null mutant lungs

To examine whether abnormal cell death accounted for the hypoplasia of *Nolz-1* null mutant lung, we assayed apoptosis using immunostaining of active Caspase-3 (AC3), an apoptosis marker. The results showed that none, at most few AC3-positive cells, were observed in wild type and KO lungs, and no apparent differences were detected at E12.5 (Figure S2A) and E13.5 (Figure S2B). Note that AC3 immunostaining was validated by positive controls in which many AC3-positive cells were present in the dorsal root ganglion (DRG) in the same sections at E12.5 and E13.5. Similar results were obtained using the terminal deoxynucleotidyl transferase-mediated dUTP nick end labeling (TUNEL) assay. Few TUNEL-positive cells were present in wild type and *Nolz-1* KO lungs without significant differences between the two genotypes at E12.5 (Figure S2C) and E13.5 (Figure S2C). TUNEL-positive cells were present in the DRG of the same sections of E12.5 embryos as a positive control. These findings suggest that *Nolz-1* null mutation does not induce abnormal apoptosis at the early stages of lung development.

### Defective cell proliferation in developing *Nolz-1* null mutant lungs

Given the hypoplasia phenotype and specific expression of Nolz-1 in the mesenchyme, *Nolz-1* null mutation might cause defective proliferation of mesenchymal cells. We then assayed cell proliferation with the BrdU incorporation assay. Pregnant mice were pulse-labeled with BrdU (50 mg/kg) for 1 h before embryo culling at E12.5 or E13.5. BrdU-positive cells were decreased by 33% and 20%, respectively, in the mutant mesenchyme at E12.5 and E13.5 compared to wild type (Figure 3A). Because branching morphogenesis was impaired in *Nolz-1* KO lungs, defective cell proliferation might occur at the distal lung buds. Double immunostaining of BrdU and Sox9, a distal epithelium marker, showed that the percentage of BrdU and Sox9 double-positive cells were, however, not different between E12.5 wild type and KO lungs (Figure S3).

**Figure 3.**
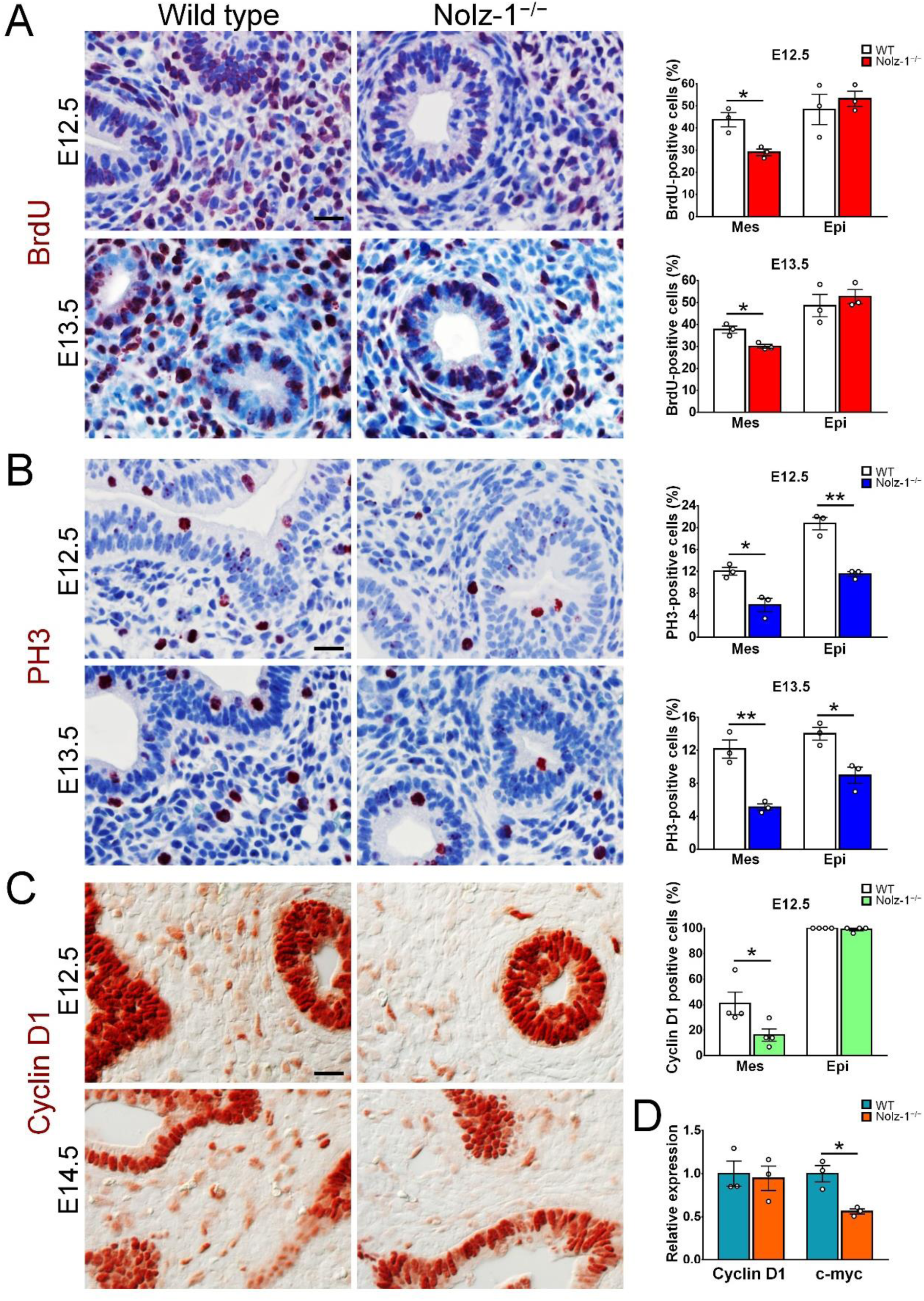
Reduced proliferating mesenchymal and epithelial cells in *Nolz-1* null mutant lungs. (A) BrdU-pulse labeling shows that BrdU-positive cells in the mesenchyme, but not in the epithelium, are decreased in E12.5 and E13.5 mutant lungs. (B) PH3-positive mitotic cells in the mesenchyme and epithelium are decreased in E12.5 and E13.5 mutant lungs. (C) Cyclin D1-positive cells in the mesenchyme, but not in the epithelium are decreased in E12.5 and E14.5 mutant lungs. (D) qRT-PCR assays show a reduction in *c-myc* mRNA in E12.5 mutant lungs. Student’s *t*-test, \**P* < 0.05, n = 3 for each group. Scale bars in A, B, C, 20 μm.

We also immunohistochemically examined the level of phosphohistoneH3 (PH3), a mitotic marker for G2/M phases (Hendzel et al., 1997; Yin et al., 2008). At E12.5, PH3-positive cells were significantly reduced by 51% and 44%, respectively, in the mesenchyme and epithelium of mutant lungs (Figure 3B). At E13.5, PH3-positive cells were decreased by 58% and 36%, respectively, in the mesenchyme and epithelium of mutant lungs (Figure 3B).

Consistent with the decreases in BrdU- and PH3-positive cells in *Nolz-1* KO lungs, cyclin D1, a G1 cyclin protein required for the progression of the G1 phase (Baldin et al., 1993), was reduced by 57% in the mesenchyme of mutant lungs at E12.5 (Figure 3C), but not in the mutant epithelium (Figure 3C). A similar reduction in cyclin D1 was found in E14.5 mutant mesenchyme. Moreover, qRT-PCR showed that c-Myc mRNA, a key regulator of proliferation (Kioussi et al., 2002), was reduced by 43% in E12.5 mutant lungs compared to wild type lungs (Figure 3D). Note that qRT-PCR show no changes in cyclin D1 mRNA in mutant lungs (Figure 3D). As whole lung tissue was collected for qRT-PCR, the predominant expression of cyclin D1 in the epithelium could have overshadowed the low level of cyclin D1 in the mesenchyme.

Taken together, the reductions in BrdU-, PH3- and cyclin D1-positive cells and c-Myc mRNA in KO lungs suggest that Nolz-1 not only cell-autonomously regulates the proliferation of mesenchymal cells, but also non-cell autonomously affects epithelial cells at the mitotic phase in developing lungs.

### Differentiation of smooth muscle and vascular endothelial cells in *Nolz-1* null mutant lungs

The progenitor cells in lung mesenchyme give rise to different cell types, including smooth muscle cells and vascular endothelial cells. Immunostaining showed that smooth muscle actin (SMA) expression was transiently decreased in *Nolz-1* KO lungs at E11.5 (Figure S4A). The SMA expression pattern was, however, not significantly altered at E12.5, E14.5, E16.5 and E18.5 in mutant lungs compared to that of wild type lungs (Figure S4A). The expressions of *SMA* and *SM22α* mRNAs, progenitor markers of smooth muscle cells, were not changed in E12.5 mutant lungs (Figure S4B). The expression pattern of PECAM-1/CD31, a marker of vascular endothelial cells, appeared to be not changed in E13.5, E16.5 and E18.5 mutant lungs (Figure S4C). Nor was Foxp1, which was expressed in the mesenchyme and distal parts of the epithelium (Shu et al., 2007), altered in E13.5 mutant lungs (Figure S4D).

### Abnormal differentiation of epithelial cells in *Nolz-1* null mutant lungs

Immunostaining showed that the TTF1/Nkx2.1 expression was not changed in E11.5 *Nolz-1* KO lungs (Figure 4A), but Foxp2, a marker of the distal part of epithelial cells (Shu et al., 2007), was ectopically expressed in the mesenchyme of *Nolz-1* mutant lungs. Notably, the ectopic expression of Foxp2 was domain-specific, i.e., Foxp2 was ectopically expressed in mesenchyme primarily surrounding the proximal parts of the epithelium (Figure 4B), suggesting that Nolz-1 suppresses *Foxp2* in the mesenchyme. Sox9 is also known to be expressed in the distal epithelial progenitors and deletion of Sox9 affects lung branching (Rockich et al., 2013). However, the Sox9 expression pattern appeared unchanged in mutant lungs at E12.5 and E14.5. The density of Sox9 positive cells was not altered in E12.5 mutant lungs (Figure 4C).

**Figure 4.**
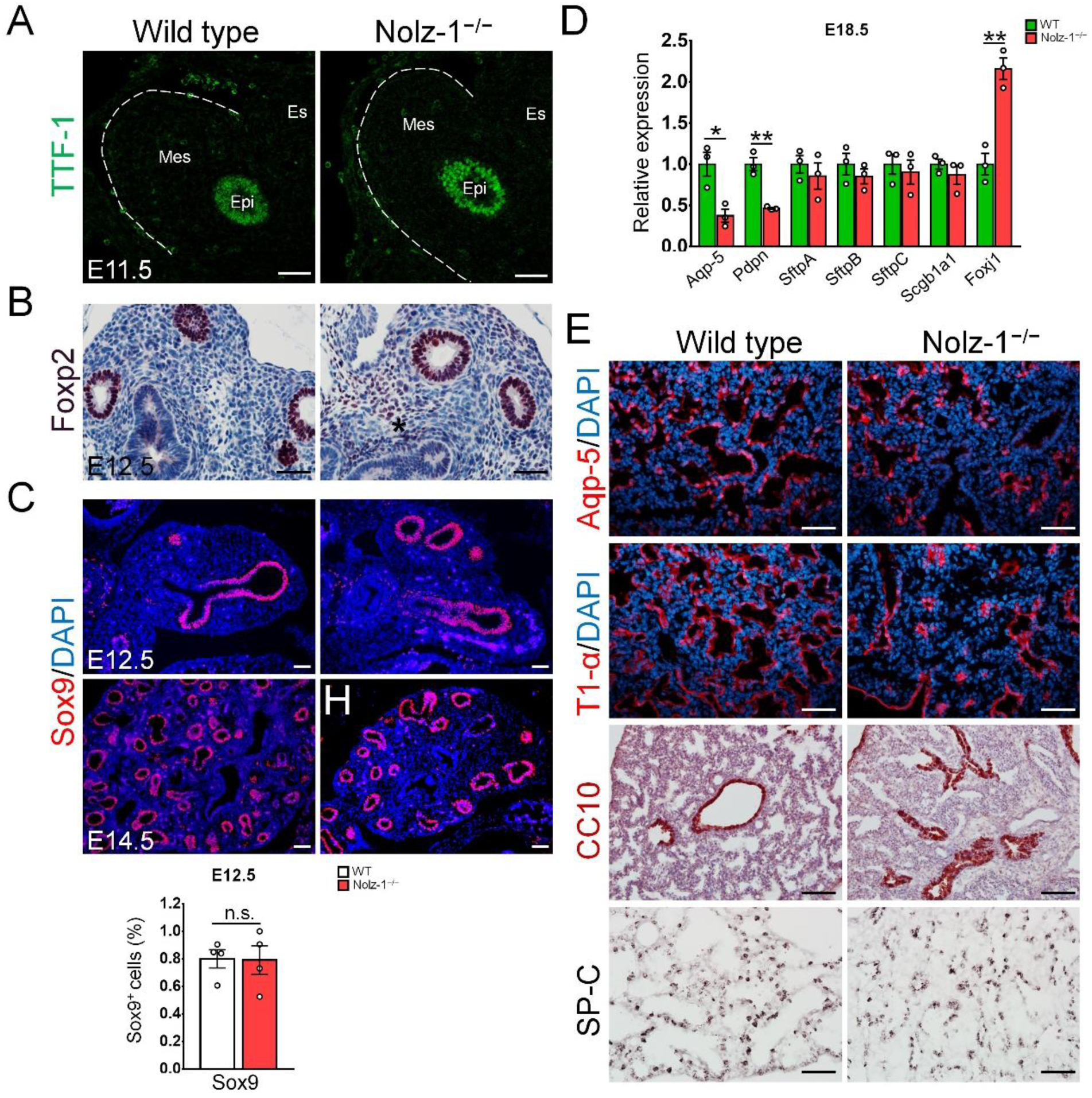
The effects of *Nolz-1* null mutation on differentiation of epithelial cells in developing mouse lungs. (A) TTF-1-positive epithelial cells are normal in E11.5 *Nolz-1* mutant lungs. (B) Foxp2 is expressed in the distal parts of the epithelium of E12.5 wild type lungs, but is ectopically expressed in the mesenchyme of E12.5 *Nolz-1* mutant lung. The ectopic region is indicated by the asterisk. (C) Sox9-positive distal tip epithelial cells were normal in E12.5 and E14.5 *Nolz-1* mutant lungs compared to the wild type. Student’s *t*-test, n.s. not significant, n = 4. (D) The AEC1 markers, *Aqp5* and *Pdpn* mRNAs, are decreased in E18.5 mutant lungs. The ciliated cell marker of*Foxj1* mRNA is increased in the mutant lungs. Student’s *t*-test, \**P* < 0.05, \**P* < 0.01, n = 3. (E) Aquaporin 5-positive cells and T1-alpha-positive cells are decreased in E18.5 *Nolz-1* mutant lungs. Many collapsed epithelia containing CC10-positive Clara cells ar present in E18.5 mutant lungs. No apparent changes in SP-C-positive AEC2 cells are found in E18.5 mutant lungs. Epi: epithelium; Mes: mesenchyme; Es: esophagus. Scale bars in A, B, C, E, 50 μm.

We performed qRT-PCR to assay different differentiated cell types in E18.5 lung epithelium using the following epithelial cell markers: alveolar epithelial type I cell (AEC1) cells: aquaporin 5 (*Aqp5*); T1-alpha (*Pdpn*), AEC2 cells: Sp-A (*SftpA*), Sp-B (*SftpB*) and Sp-C (*SftpC*); Clara cells: CC10 (*Scgb1a1*); ciliated cells: *Foxj1*. The results showed that *Aqp5* and *Pdpn* mRNAs were decreased in E18.5 mutant lungs, whereas *Foxj1* mRNA was increased in mutant lungs. *SftpA*, *SftpB*, *SftpC* and *Scgb1a1* mRNAs were not changed in mutant lungs (Figure 4D). Consistent with the qRT-PCR results, the immunostaining showed Aqp5 and T1-alpha expressions were decreased in E18.5 *Nolz-1* KO lungs. Collapsed CC10-positive epithelia were evident in E18.5 mutant lungs (Figure 4E). SP-C-positive AEC2 cells appeared to be no affected in E18.5 mutant lungs (Figure 4E). These results suggest that Nolz-1 may non-cell autonomously regulate the differentiation of epithelial cells.

### Quantitative RT-PCR profiling of morphogenic molecules in *Nolz-1* mutant lungs

Previous studies have shown the importance of morphogenic molecules for lung development, including Wnts, Fgfs, Pdgfs, Shh, Bmps and Tgf-β2 (Morrisey and Hogan, 2010; Warburton et al., 2000; Zepp and Morrisey, 2019). Hypoplasia of mouse lung has previously been documented in Shh, Fgf9, Fgf10 and Wnt2 null mutant mice, and defective cell proliferation of epithelial and mesenchymal cells was also found in these mutant mice (Colvin et al., 2001; Goss et al., 2009; Pepicelli et al., 1998; Sekine et al., 1999). We then examined whether these signal molecules were altered in E12.5 *Nolz-1* KO lungs with qRT-PCR. The results showed that *Wnt2*, *Lef1*, *Fgf10*, *Gli3* and *Bmp4* were decreased in mutant lungs (Figure 5A). In contrast, *Pdgfrα* and *Tgf-β2* were increased in mutant lungs (Figure 5A). The expression of *Wnt2b*, *Wnt7b*, *Axin2*, *Fgf9*, *Fgf*r2, *Pdgfc*, *Pdgfrβ*, *Shh* and *Gli2* mRNAs were not changed in mutant lungs (Figure 5A). These results suggest that Nolz-1 may regulate Wnt, Shh, Bmp, Pdgf and Tgf signaling in developing lungs.

**Figure 5.**
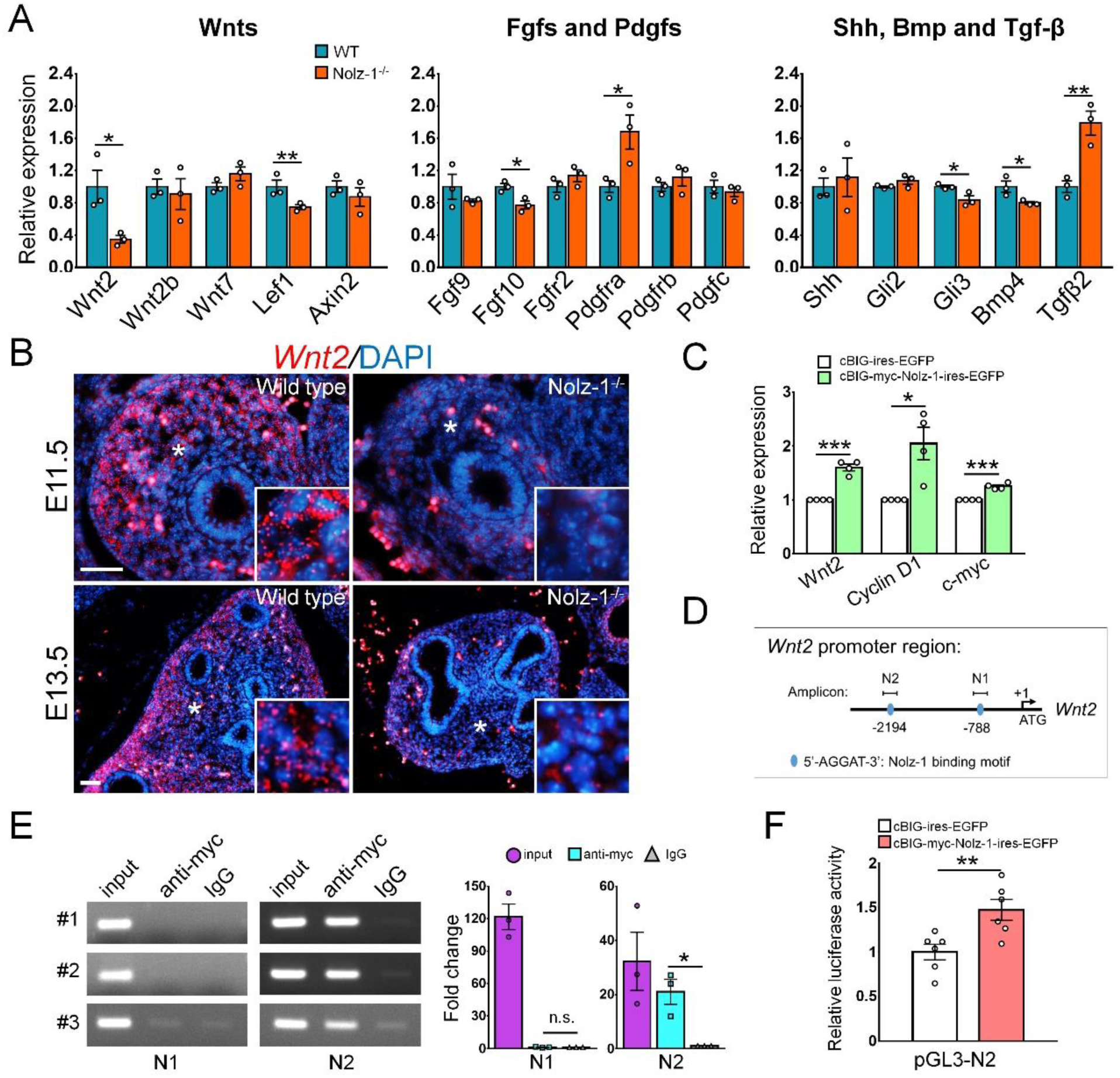
*Wnt2* is a Nolz-1 target gene in developing lungs. (A) Morphogenic molecules are screened by qRT-PCR in E12.5 *Nolz-1* mutant lungs. *Wnt2* and *Lef1* mRNA are decreased in *Nolz-1* mutant lungs. *Fgf10* is decreased and *Pdgfrα* is increased in the *Nolz-1* mutant lungs. *Gli3* and *Bmp4* are decreased and *Tgf-β2* is increased in *Nolz-1* mutant lungs. Student’s *t-*test, * *P* < 0.05, \*\**P* < 0.01, n = 3. (B) *In situ* hybridization shows that *Wnt2* mRNA is mainly expressed in the distal parts of the mesenchyme of E11.5 and E13.5 wild type lungs. *Wnt2* mRNA is markedly decreased in mutant mesenchyme. The insets show high magnification of the regions indicated by asterisks. Scale bars, 50 μm. (C) The qRT-PCR assay shows that over-expression of Nolz-1 by electroporation of pcBIG-myc-Nolz-1-ires-EGFP plasmid up-regulates *Wnt2*, *cyclinD1* and *c-myc* mRNAs in primary mesenchymal cell culture derived from E14.5 wild type mouse lungs compared to mock transfection of pcBIG-ires-EGFP control plasmids. Student’s *t-*test, * *P* < 0.05, \*\*\**P* < 0.001, n = 4. (D) The schematic drawing illustrates the locations of the putative Nolz-1 binding sites of “AGGAT” at -788 (N1 motif) and -2194 (N2 motif) in 5 ‘flânking regions of mouse *Wnt2* gene (+1: ATG translation start site). (E) The chromatin immunoprecipitation (ChIP) assay shows that a 187 bp PCR band is amplified from the N2 locus with the immunoprecipitated products using the anti-myc antibody, but not the control rabbit IgG, in E14.5 lung mesenchymal cell culture electroporated with pcBIG-myc-Nolz-1-ires-EGFP plasmids. No specific PCR band is detected from the N1 locus. Student’s *t-*test, * *P* < 0.05, n.s. not significant, n = 3. (F) The reporter gene assay showed that the luciferase activity is increased in the pGL3-N2-Luc group compared to the pGL3-Luc control group in E14.5 lung mesenchymal cells transfected with pcBIG-myc-Nolz-1-ires-EGFP plasmids. Student’s *t-*test, \*\**P* < 0.01, n = 6.

### Down-regulation of Wnt2 in the mesenchyme of *Nolz-1* mutant lungs

Previous studies have shown that Wnt2 (Wnt2a) regulates the proliferation of mesenchymal cells in developing mouse lungs (Goss et al., 2009). Because *Wnt2* expression was drastically reduced by 66% in *Nolz-1* mutant lungs, we asked whether Wnt2 signaling was involved in the regulation of cell proliferation in *Nolz-1* mutant lungs. We validated *the* reduction in *Nolz-1* mutant lungs by in situ hybridization. *Wnt2* mRNA was mainly expressed in the distal parts of mesenchyme of E11.5 and E13.5 wild type lungs (Figure 5B). Consistent with the result of qRT-PCR. 5a), *Wnt2* mRNA was markedly decreased in the mutant mesenchyme.

### Over-expression of *Nolz-1* upregulates *Wnt2*, *cyclinD1* and *c-myc* in the mesenchymal culture of developing lungs

The loss-of-function study showed that Wnt2 expression was decreased in the mesenchyme of *Nolz-1* mutant lungs. We then asked whether Nolz-1 was sufficient to induce *Wnt2* in lung mesenchymal cells. We cultivated primary mesenchymal cells from E14.5 wild type mouse lungs. Cultured cells were electroporated with pcBIG-myc-Nolz-1-ires-EGFP and pcBIG-ires-EGFP control plasmids. Electroporated cells were cultivated for 48 hr and were then harvested for the qRT-PCR assay (Figure S5). Nolz-1 over-expression markedly increased *Wnt2* mRNA by 60% compared to the mock control pcBIG-ires-EGFP (Figure 5C). It also significantly increased *cyclin D1* and *c-myc* mRNAs by 104% and 24%, respectively, compared to mock controls in mesenchymal cells (Figure 5C).

Collectively, the loss-of-function and gain-of-function studies suggest that Nolz-1 is sufficient and required for the expression of *Wnt2*, *cyclin D1* and *c-myc* in mesenchymal cells of developing lungs.

### Wnt2 is a direct target gene of *Nolz-1* in developing lungs

We further asked whether Nolz-1 directly regulated *Wnt2* in lung mesenchymal cells. Previous studies have identified the DNA sequence “5’-AGGAT-3’” as Nolz-1 putative binding motif in *zebrafish* (Runko and Sagerstrom, 2003; Siggers et al., 2014). We searched the 5’ flanking region of the *Wnt2* mouse gene and found two “AGGAT” motifs at -788 (N1) and -2194 (N2) in the 5’ flanking regions (+1: translation start site) (Figure 5D). We electroporated pcBIG-myc-Nolz-1-ires-EGFP in the primary culture of mesenchymal cells of E14.5 lungs. Two days after electroporation, cultured cells were harvested for the chromatin immunoprecipitation (ChIP) assay. A PCR band with the expected size of 187 bp was amplified from the N2 locus with the anti-myc epitope-tag antibody, but not with the control rabbit IgG (Figure 5E). No specific PCR band was detected at the N1 locus (Figure 5E).

Next, we cloned the 187 bp DNA containing the N2 motif into the pGL3-basic luciferase (Luc) reporter plasmid. The pGL3-N2-Luc or its control pGL3-Luc plasmid was co-transfected with pcBIG-myc-Nolz-1-ires-EGFP or its mock control pcBIG-ires-EGFP plasmid in E14.5 primary culture of mesenchymal cells. Cells were cultured for 48 hr before harvest for the reporter gene assay. Luc activity was increased by 47% in pGL3-N2-Luc compared to pGL3-Luc control groups (Figure 5F). These results suggest that Nolz-1 can bind to the “AGGAT” of the N2 motif in the promoter region of *Wnt2* to transcriptionally activate the expression of Wnt2 in lung mesenchymal cells.

### Nolz-1 regulates canonical Wnt/β-catenin signaling in developing lungs

Previous studies have demonstrated that Wnt2/2b regulates lung development through the canonical Wnt/β-catenin pathway (Goss et al., 2011; Goss et al., 2009). Given that *Wnt2* was a target gene of Nolz-1, we asked whether Nolz-1 was involved in the Wnt/β-catenin signal pathway. We intercrossed *Nolz-1+/−* mice with *BATGAL* or *TOPGAL* Wnt/β-catenin transgenic reporter mice to generate *Nolz-1−/−;BATGAL* or *Nolz-1−/−;TOPGAL* mice. LacZ staining in the whole-mount showed a reduction in LacZ signals in *Nolz-1−/−;BATGAL* and *Nolz-1−/−;TOPGAL* lung. In *BATGAL* lungs, LacZ signals were expressed in epithelial columns and the mesenchyme. LacZ-positive signals were decreased in both the epithelium and mesenchyme of *Nolz-1−/−;BATGAL* mice (Figure 6A). In *TOPGAL* lungs, LacZ signals were restricted only to epithelial columns. LacZ signals were reduced in epithelial columns of *Nolz-1−/−;TOPGAL* mutant lungs (Figure 6B). Furthermore, immunostaining of β-catenin showed decreased β-catenin immunoreactivity in E13.5 *Nolz-1 null* mutant mesenchyme (Figure 6C). Axin2 immunoreactivity was reduced in the distal parts of E13.5 *Nolz-1* mutant mesenchyme (Figure 6D). However, Axin2 expression appeared normal in the mesenchymal regions surrounding the airways of mutant lungs (Figure 6D).

**Figure 6.**
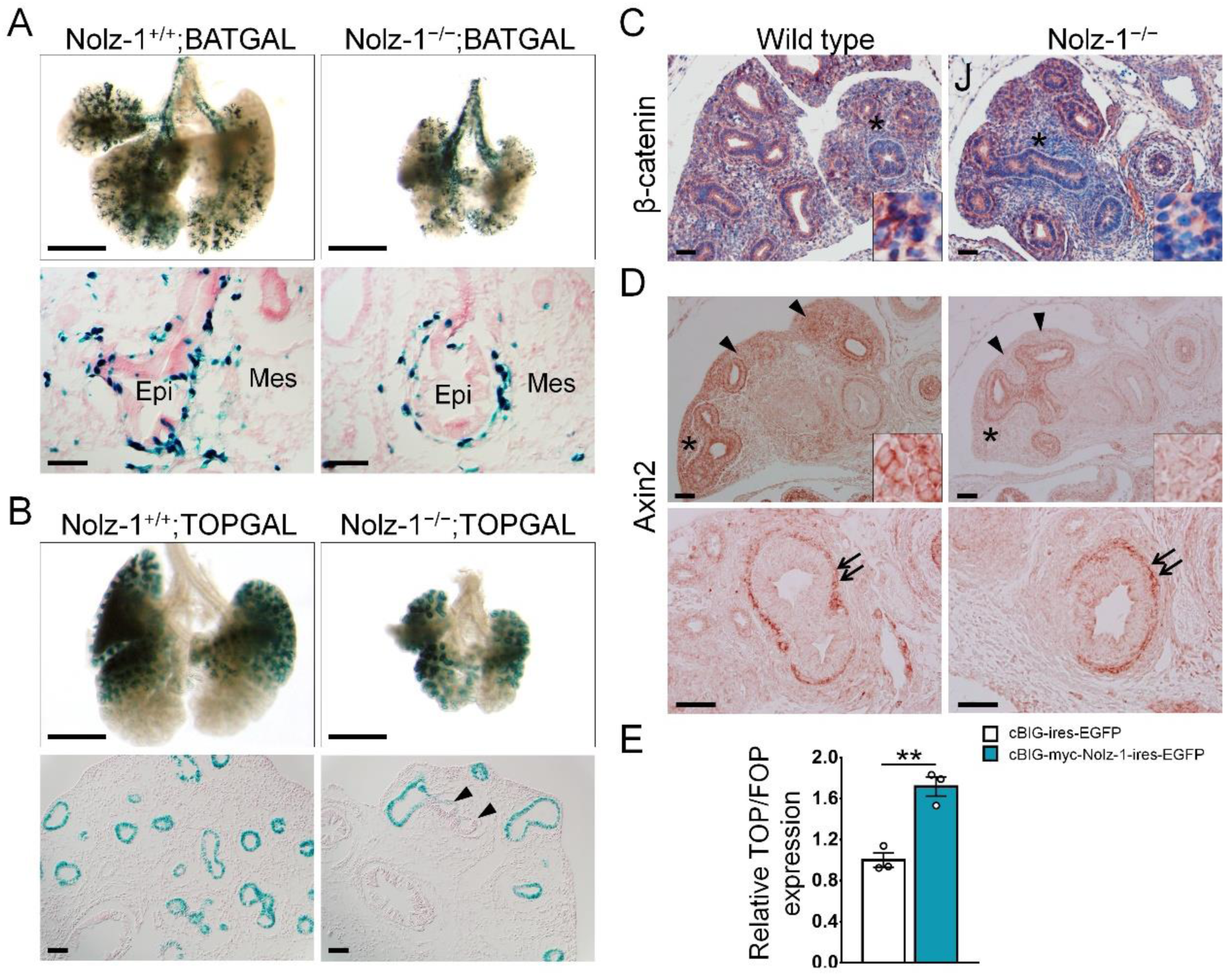
Nolz-1 regulates canonical Wnt/β-catenin signaling in developing lungs. (A) LacZ staining of E14.5 *Nolz-1+/+;BATGAL* (control) and *Nolz-1−/−;BATGAL* mutant lungs. Decreased lacZ expression in the developing epithelium and mesenchyme was observed in the mutant lungs. (B) Whole-mount LacZ staining of E14.5 *Nolz-1+/+;TOPGAL* (control) and *Nolz-1−/−;TOPGAL* mutant lungs shows decreased lacZ expression in the lung epithelium. Arrowheads indicate epithelial cells without LacZ signals in the *Nolz-1* mutant lung. (C) β-catenin immunoreactivity is decreased in the E13.5 *Nolz-1* mutant mesenchyme compared to that of wild type. The insets show high magnification of the regions indicated by asterisks. (D) Axin2 immunoreactivity is reduced in the E13.5 *Nolz-1* mutant lung mesenchyme (arrowheads). The insets show high magnification of the regions indicated by asterisks. Axin2 immunoreactivity appears to be not altered in the regions around the airway (arrows) of the *Nolz-1* mutant lungs. (E) Over-expression of Nolz-1 increases the ratio of TOPFLASH/FOPFLASH activity in lung mesenchymal cells. Student’s *t*-test, \*\**P* < 0.01, n = 3. Scale bars in A, B, 500 μm; C, D, 50 μm.

We next examined whether Nolz-1 activated Wnt/β-catenin signaling with the TOPFlash/FOPFlash assay in the mesenchymal culture of lungs. TOPFlash and FOPFlash are luciferase reporters that contain normal and mutant TCF-binding sites, respectively. The ratio of TOPFlash/FOPFlash represents the specific activity of Wnt/β-catenin signaling. Over-expression of Nolz-1 significantly increased the TOPFlash/FOPFlash ratio by 71% compared to the mock control (Figure 6E).

### Exogenous Wnt2 rescues epithelial branches and proliferation of mesenchymal cells in *Nolz-1* mutant lungs

The above findings suggest that Nolz-1 may regulate mesenchymal cell proliferation through the canonical Wnt2/β-catenin signal pathway. Since deletion of *Nolz-1* resulted in the reductions in Wnt2 expression and proliferation of mesenchymal cells, we asked whether exogenously applied Wnt2 protein could causally rescue defective proliferation of mesenchymal cells in *Nolz-1* KO lungs. We cultivated 12.5 lung explants with recombinant human Wnt2 protein (rWnt2, 150 ng/ml) for 48 hr. Lung explant culture was pulse-labeled with BrdU for 1 hr before tissue harvest. The number of epithelial branches at 48 hr after culture was normalized with that at the beginning of culture (0 hr). In wild type lung explants, there was no difference between vehicle and rWnt2 treatments in epithelial branches (Figure 7A). In *Nolz-1* KO lung explants, rWnt2 treatments markedly increased epithelial branches compared to the vehicle group (Figure 7A). Treatments with rWnt2 also significantly increased proliferating BrdU-positive cells by 61% compared to the vehicle group (Figure 7B). In wild type lung explants, rWnt2 treatments did not increase BrdU-positive cells in the mesenchyme (Figure 7B). The endogenous Wnt2 protein in wild type lung explants may have diminished the effects of exogenous Wnt2. *Nolz-1* deletion decreased the endogenous Wnt2 level, which may allow exogenous rWnt2 to promote the proliferation of mesenchymal cells and the growth of epithelial branches in mutant lung explants. The rescue experiment supports the causality that Nolz-1 regulates mesenchymal cells and epithelial branches through Wnt2a signaling.

**Figure 7.**
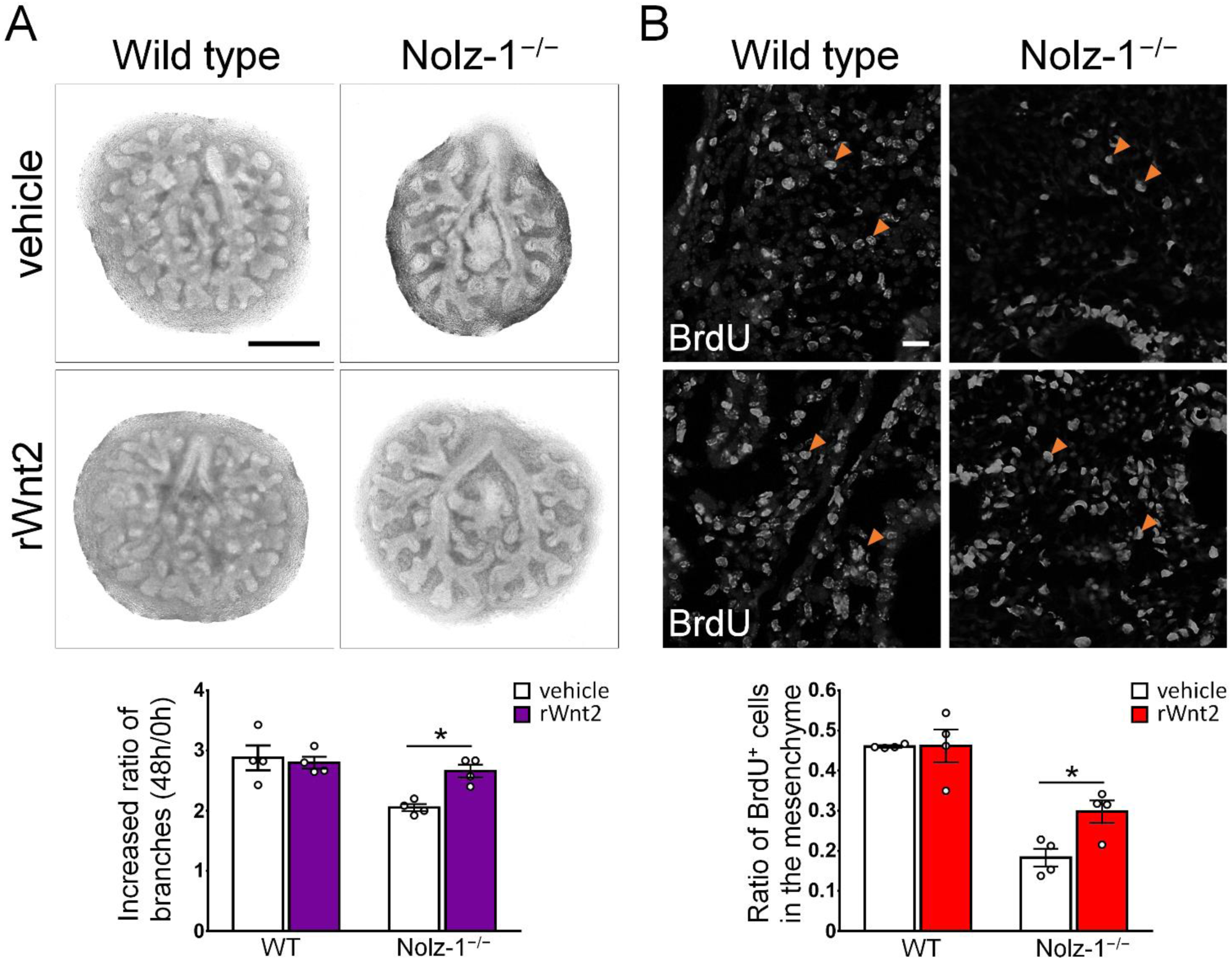
Exogenous rWnt2 rescues mesenchymal cell proliferation and epithelial branching in the culture of *Nolz-1* null mutant lungs. (A) Treatments with rWnt2 (150 ng/ml) increase the number of epithelial branches in the *Nolz-1* mutant lung culture compared to vehicle control. The number of epithelial branches at 48 hr after cultivation is normalized with that at the beginning of culture (0 hr). Student’s *t*-test, \**P* < 0.05, n = 4. (B) Treatments with rWnt2 significantly increase BrdU-positive cells (arrowheads) in mutant lung culture. Student’s *t*-test, \**P* < 0.05, n = 4. Scale bars in A, 500 μm; B, 20 μm.

### Fgf9 regulated *Nolz-1* expression in developing lungs

Finally, we attempted to identify upstream signaling molecules that regulate *Nolz-1* expression in developing lungs. We focused on Fgf9 and Fgf10, because Fgf9 and Fgf10 promote the proliferation of mesenchymal cells in the developing lungs (del Moral et al., 2006; Jean et al., 2008). Wild type lung explant culture was treated with human recombinant Fgf 9 (rFgf9) or rFgf10 for 48 hr. Treatments with rFgf10 did not alter the epithelial structure, nor did they change *Nolz-1* and *Wnt2* mRNAs (Figure 8A). By contrast, rFgf9 treatments not only resulted in enlarged epithelia, but also significantly increased *Nolz-1*, *Wnt2*, *Lef1* mRNAs compared to vehicle treatments (Figure 8B). Consistently, Western blotting showed a marked increase in Nolz-1 protein with rFgf9 compared to the vehicle group (Figure 8B). These findings suggest that Fgf9 signaling acts upstream of Nolz-1 in developing lungs.

**Figure 8.**
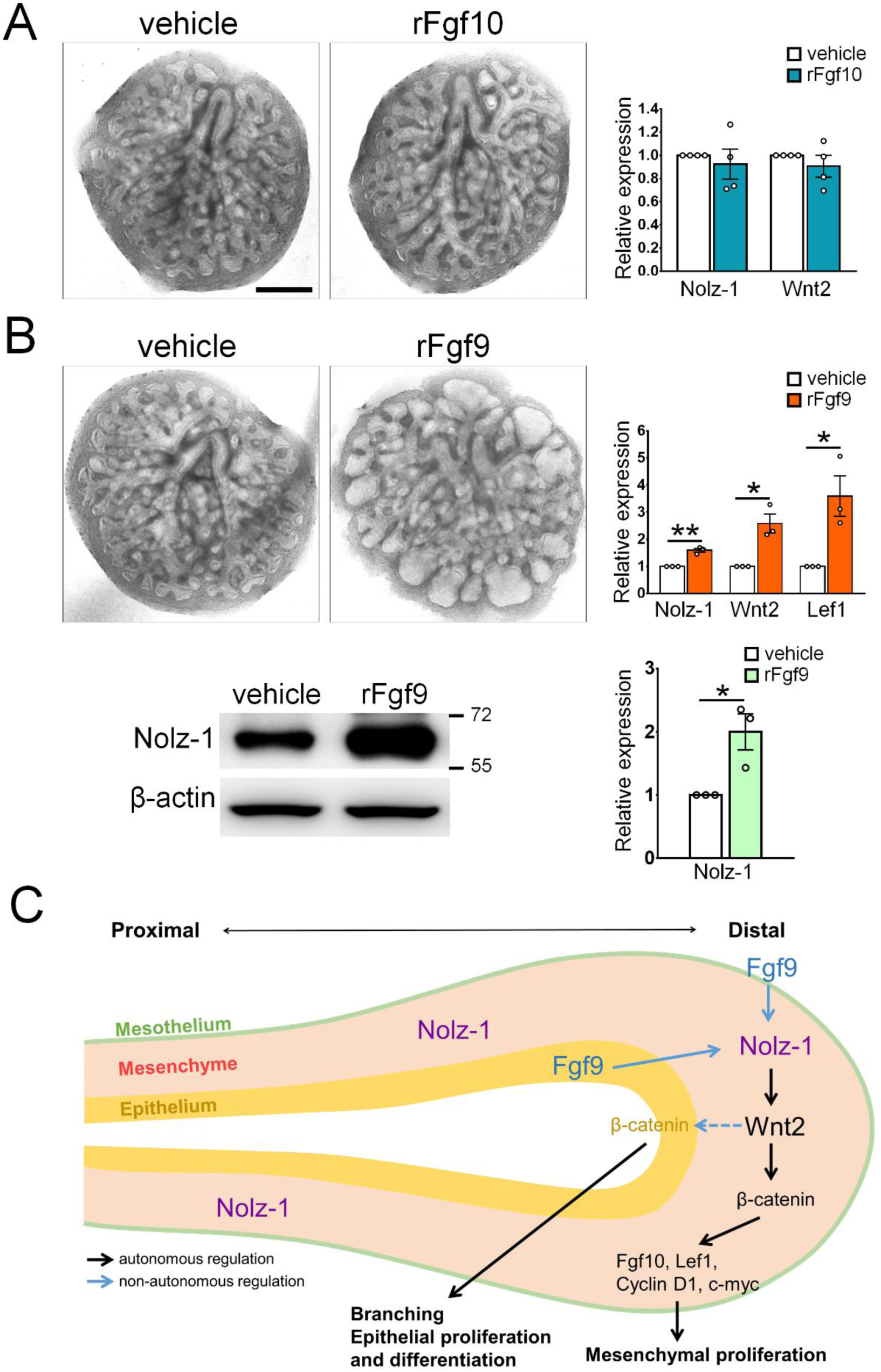
Fgf9 promotes *Nolz-1* expression in developing lungs and working hypothesis. (A) Treatment with rFgf10 (200 n /ml) of wild type explant lung culture for 48 hr. qRT-PCR shows that *Nolz-1* and *Wnt2* are not changed in the rFgf10 treated group compared to the vehicle control. Student’s *t*-test, *P* > 0.05, n = 4. Scale bar, 500 μm. (B) Treatment with rFgf9 (200 ng/ml) results in enlarged epithelia in wild-type explant lungs cultured for 48 hr. The qRT-PCR shows that *Nolz-1*, *Wnt2* and *Lef1* mRNAs are increased in rFgf9 treated group in wild type lungs. Student’s *t*-test, \**P* < 0.05, \*\**P* < 0.01, n = 4. Western blotting showed that rFgf9 treatment increases Nolz-1 protein by 99% in wild type lung culture compared to the vehicle-treated group. Student’s *t*-test, \**P* < 0.05, n = 3. (C) Working hypothesis. Nolz-1 controls the proliferation of mesenchymal cells and the growth of epithelial branches through the regulation of Wnt2 signaling in the early stages of the development of the lungs. In the late stages of development, Nolz-1 acts non-cell autonomously to regulate the development of epithelial cells through Wnt2 signaling. Fgf9 acts upstream to regulate *Nolz-1* expression in developing lungs.

## Discussion

We have identified Nolz-1 as a mesenchyme-specific transcriptional regulator that plays an essential role in lung morphogenesis. Nolz-1 regulated mesenchymal cell proliferation and the growth of epithelial branches. Loss-of-function and gain-of-function studies identified Wnt2 as a candidate molecule for mediating Nolz-1 function in developing lungs. Furthermore, we showed that Fgf9 is an upstream regulator of *Nolz-1*. We propose that the Fgf9-Nolz-1-Wnt2 signaling axis plays an important role in controlling lung morphogenesis (Figure 8C).

### Altered morphogenic molecules in Nolz-1 mutant lungs

Previous studies have shown that several morphogens, including Shh, Fgfs, Wnts and BMP proteins can regulate the proliferation of mesenchymal cells in developing lungs (Morrisey et al., 2013). Epithelial Shh promotes cell proliferation and BMP4 expression in the mesenchyme during lung bud formation2. Null mutation of Fgf9 derived from the epithelium- and mesothelium causes defective cell proliferation in the lung mesenchyme (White et al., 2007; White et al., 2006). Epithelium-derived Wnt7b promotes cell proliferation in the mesoderm of the lung through Axin2 and Lef1 (Rajagopal et al., 2008; Shu et al., 2002). The Wnt2 null mutation results in significant lung hypoplasia due to decreased cell proliferation in both the epithelial and mesenchymal compartments (Goss et al., 2009). In most cases, these secreted morphogenic molecules regulate cell proliferation in both compartments of the epithelium and mesenchyme.

We have identified Nolz-1 as an important transcription factor that regulates cell proliferation in the mesenchyme. Because Nolz-1 was expressed in the lung primordium as early as E11.5, at which time lung morphogenesis begins to occur, Nolz-1 may regulate morphogenic molecules, which in turn control cell proliferation in the lung mesenchyme. We examined Shh, Fgfs, Wnts and BMP signaling molecules to determine whether Nolz-1 regulated cell proliferation through these morphogenic molecules. The qRT-PCR assay showed that the expressions of *Wnt2* and *Fgf10* were decreased in E12.5 mutant lungs (Figure 5A). The concurrent decreases in *Wnt2* and *Fgf10* were interesting, because *Wnt2* null mutation has been shown to decrease *Fgf10* in developing mouse lungs (Goss et al., 2011). Therefore, deletion of *Nolz-1* leads to *Wnt2* reduction, which may in turn down-regulate *Fgf10* in the *Nolz-1* mutant lungs.

### Nolz-1 as a mesenchyme-specific transcriptional regulator

Previous studies have identified several transcription factors that are specifically expressed in the lung mesenchyme, including Foxf1, Tbx4/5 and Hox5 (Cardoso and Lu, 2006; Morrisey and Hogan, 2010). Deletion of Foxf1 inhibits mesenchymal cell proliferation and delays lung branching with altered expression of Wnt2, Hoxb7 and Wif genes (Ustiyan et al., 2018). Tbx4/5 mutants exhibit defective lung branching and down-regulation of the mesenchymal markers of Wnt2 and Fgf10 and the epithelial markers of Bmp4 and Spry2 (Arora et al., 2012). Hoxa5;Hoxb5;Hoxc5 triple-mutant embryos show deficits in lung branching and proximal-distal patterning. Deletion of Hox5 leads to loss of Wnt2/2b expression in the distal lung mesenchyme and reduction of downstream Wnt2 signaling molecules of Lef1 and Axin2 in the mesenchyme and Bmp4 in the epithelium (Hrycaj et al., 2015).

It is of particular interest that *Nolz-1* KO lungs share some phenotypes, including decreased cell proliferation in lung buds and down-regulation of mesenchymal Wnt2 signaling, with Foxf1, Tbx4/5 and Hox5 KO lungs, suggesting that Nolz-1 may interact with these mesenchyme-specific transcription factors in the regulation of lung development. Indeed, we found that the Hoxa5 and Hoxb5 expressions were reduced in *Nolz-1* KO lungs (S.-Y. Chen and F.-C. Liu, unpublished observations).

### Nolz-1-Wnt2 signaling axis in the control of lung morphogenesis

Our ChIP and reporter gene data suggest that Nolz-1 transcriptionally regulates *Wnt2* in developing lungs. We postulate that Nolz-1 may regulate mesenchymal cell proliferation and epithelial branching through Wnt2 signaling. This hypothesis is supported by the rescue experiment in which the exogenous rWnt2 protein could partially rescue defective cell proliferation and epithelial branching in mutant lung culture (Figure 7). Consistent with this hypothesis, the lung phenotypes of *Nolz-1* KO lung phenotypes were similar to that of *Wnt2* (*Wnt2a*) KO mice (Goss et al., 2009). *Wnt2* deletion induces lung hypoplasia in which reduced cell proliferation occurs in both the epithelial and mesenchymal compartments. In *Wnt2/2b* double knockout mice, Nkx2.1-positive lung endoderm progenitors are lost and the trachea and epithelial branches fail to develop in the mutant lungs. Note that deletion of *Nolz-1* selectively decreased *Wnt2* but not *Wnt2b* in developing lungs.

Deletion of *Wnt2* also results in decreased SMA expression in smooth muscle cells, and Wnt2 can promote the development of smooth muscle of the lung airway through myocardin/Mrtf-B and Fgf10 (Goss et al., 2011). Notably, a transient decrease in SMA expression was detected in *Nolz-1* knockout lungs at E11.5, although no significant changes in SMA immunoreactivity were found in later stages (Figure S4A). Given that Nolz-1 positively regulates *Wnt2*, Nolz-1 may act through Wnt2 signaling to regulate not only cell proliferation but also smooth muscle cell differentiation in the lung mesenchyme. Our study also showed that Nolz-1 was engaged in canonical Wnt/β-catenin signaling in lung morphogenesis. The data of *BATGAL* and *TOPGAL* reporter mice and the TOPFlash/FOPFlash assay indicated that Nolz-1 is engaged in canonical Wnt/β-catenin signaling in developing lungs. Wnt2 is known to act through the canonical Wnt/β-catenin pathway in lung mesenchymal cells (Goss et al., 2011; Yin et al., 2008). *Nolz-1* KO not only decreased Wnt2, but also reduced downstream target genes of canonical Wnt/β-catenin signaling, including *Lef1*, *c-myc,* and *cyclin D1* (De Langhe et al., 2008; Juan et al., 2014; Shtutman et al., 1999; Tetsu and McCormick, 1999; Zhang et al., 2012).

### Fgf9-Nolz-1-Wnt2 signaling in the regulation of lung morphogenesis

Because Nolz-1 was expressed as early as E11.5 when cell proliferation was activated by morphogen molecules, the earliest time point examined in the present study, the proliferative effects of morphogenic molecules may be mediated by Nolz-1 signaling. Fgf9 is required for the proliferation of mesenchymal and epithelial cells (del Moral et al., 2006; White et al., 2006). Previous study has shown reductions in Wnt2 and cell proliferation in Fgf9 KO lungs. Transgenic over-expression of Fgf9 in the epithelium increases Wnt2 in the lung mesenchyme (Yin et al., 2008). Because Nolz-1 is required for *Wnt2* expression and cell proliferation in the lung mesenchyme, we postulate that Fgf9 acts upstream of *Nolz-1*. Consistent with this hypothesis, treatments with rFgf9 significantly induced *Nolz-1*, *Wnt2,* and *Lef1* in E12.5 lung explant culture (Figure 8B). In addition to Fgf9, Fgf10 appears to act upstream of Wnt2, as Wnt2 is decreased in the lung primordium of Fgf10 KO mice (Sekine et al., 1999). However, treatments with Fgf10 did not up-regulate *Nolz-1* in the E13.5 lung explant culture (Figure 8A). Therefore, we propose that Fgf9 secreted from the epithelium and mesothelium (White et al., 2006) may regulate *Nolz-1* expression in developing lung mesenchyme. Fgf9-Nolz-1-Wnt2 signaling represents a novel axis in the regulation of lung morphogenesis (Figure 8C).

### A potential role of Nolz-1 in epithelium-mesenchyme interaction

The developing mouse lung contains two compartments, the epithelium and mesenchyme. The epithelium of the developing lung buds is derived from the endoderm, whereas the mesenchyme surrounding the buds is derived from the mesoderm at E9.5-E10 in mouse embryogenesis (Maeda et al., 2007; Morrisey and Hogan, 2010). Transplantation experiments have shown that cross-talk signaling between the epithelium and mesenchyme is important for branching morphogenesis and cytodifferentiation (Alescio and Cassini, 1962; Sakakura et al., 1976). The signal molecules secreted from the mesenchyme regulate the growth and differentiation of epithelial cells and vice versa (Zepp and Morrisey, 2019). For instance, Fgf10 released from the distal mesoderm acts on the epithelium through Fgfr2IIIb to promote cell proliferation and bud morphogenesis (De Moerlooze et al., 2000). Fgf9 is produced by the lung mesothelium and epithelium, and Fgf9 signaling through mesenchymal Fgfr1c and Fgfr2c to maintain mesenchymal cell proliferation (Colvin et al., 2001; White et al., 2007). Wnt7b is expressed in the epithelium to promote cell proliferation in the mesoderm through Axin2 and Lef1 (Rajagopal et al., 2008; Shu et al., 2002) and the epithelial Shh promotes cell proliferation and Bmp4 expression in the mesenchyme2. Induction of epithelial branching also requires the interaction of epithelial cells with mesenchymal cells. The co-culture experiment demonstrates that the extent of lung bud induction depends on the amount of co-cultured mesenchyme (Masters, 1976).

Although Nolz-1 is specifically expressed in the compartment of mesenchymal cells, defective epithelial cell development of epithelial cells was observed in *Nolz-1* KO lungs, e.g., Aquaporin 5 and T1-alpha, markers of AEC1, were decreased, but *Foxj1*, a marker of ciliated cells, was increased in *Nolz-1* mutant lungs. Foxp2, a marker of the distal part of epithelial cells (Shu et al., 2007), was ectopically expressed in the mesenchyme of E12.5 *Nolz-1* KO lungs. The non-cell-autonomous effects of *Nolz-1* mutation may result from defective signal transduction across the mesenchymal and epithelial compartments. As early as E12.5, a reduction in epithelial proliferation was found in *Nolz-1* mutant lungs. It is of interest that defective epithelial proliferation also occurs in Wnt2 and Fgf10 mutant mice (Goss et al., 2011; Goss et al., 2009; Sala et al., 2011). Because deletion of *Nolz-1* down-regulated *Wnt2* and *Fgf10*, it is possible that the comprised Wnt2 and Fgf10 signaling in the mesenchymal compartment may non-cell autonomously affect epithelial proliferation in *Nolz-1* mutant lungs. This possibility awaits to be examined in future studies.

### Pathological role of *NOLZ-1/ZNF503* in tumorigenesis

Our study shows that Nolz-1/Znf503 regulates mesenchymal cell proliferation in lung morphogenesis. The ability of Nolz-1 to promote cell proliferation suggests a pathological role for Nolz-1 in tumorigenesis. Indeed, previous studies have shown that NOLZ-1/ZNF503/zeppo2 promotes the proliferation and invasion of mammary epithelial cells. ZNF503/zeppo2 enhances aggressive breast cancer progression by repression of GATA3 (Shahi et al., 2015; Shahi et al., 2017) and promotes the migration, invasion and EMT process of hepatocellular carcinoma cells by suppressing GATA3 (Yin et al., 2019). ZNF503 is also involved in lung cancer.

*ZNF503* mRNA level is up-regulated in non-small cell lung cancer (NSCLC). Furthermore, microRNA and long noncoding RNA (lncRNA) can regulate tumorigenesis through ZNF503, e.g., *miR-340-5p* acts through ZNF503 to inhibit proliferation and invasion in an NSCLC cell line (Lu and Zhang, 2019). The TTN-AS1 lncRNA promotes the progression of NSCLC via the *miR-491-5p/ZNF503* axis (Qi and Li, 2020). Therefore, Nolz-1 appears to play a pathogenic role in the regulation of abnormal cell proliferation in tumorigenesis.

### Evolutionary conservation of Nolz-1 in the developmental control of respiratory organs

It is of interest that the mouse *Nolz-1* gene is one of the evolutionarily conserved genes that have high degrees of conservation of the 5’ flanking promoter regions in different species (Chang et al., 2013; Chang et al., 2011). It implies that the expression pattern of Nolz-1 in different organs may be conserved between different species. Consistent with this hypothesis, the previous study has shown that the Drosophila homologs of *Nolz-1*, *Elb/noc*, are essential for the morphogenesis of the dorsal tracheal branches during development (Dorfman et al., 2002). Elb is expressed specifically in the dorsal branches of the trachea. *The Elb* mutation induces abnormal migration of the dorsal branches, whereas the misexpression of Elb in the trachea causes loss of the visceral branch and increased dorsal branch. As a mammalian member of the NET family, Nolz-1 plays an essential role in lung morphogenesis. Therefore, *Nolz-1* and its homologues are part of transcriptional regulator toolkits that are essential for the development of respiratory organs.

## Materials and Methods

### Animals

The Wnt/β-catenin transgenic reporter mice *BATGAL* mice, (Stock No: 005317) and *TOPGAL* mice (Stock No: 004623) were obtained from The Jackson Laboratory. *Nolz-1+/-* mice (Chen et al., 2020) and the *Wnt/β-catenin* transgenic reporter mice were housed in a 12 hr light-dark cycle room at the Animal Center. The protocol of animal use was approved by the Institutional Animal Care and Use Board and was in accordance with the NIH Guide for the Care and Use of Laboratory Animals. Embryonic day (E) 0.5 was defined as the day of positivity of mating plugs.

### PCR genotyping of transgenic mice

The genotypes of transgenic mice were identified by polymerase chain reaction (PCR) with genomic DNA. Genomic DNA was prepared according to the method described at The Jackson Laboratory (http://jaxmice.jax.org). Briefly, the toes or tails of mice were lysed in NaOH buffer (25 mM NaOH, 0.2 mM EDTA) and heated to 98°C for 30 min and then cooled down to room temperature. An equal volume of 40 mM Tris-HCl (pH 5.5) was added to the lyse buffer to neutralize NaOH. After spinning down briefly, the genomic DNA was collected and used as templates for PCR reaction with pairs of genotype-specific specific primers in the followings: Nolz-1 floxed-Forward (5’-CCAAT GTGGA AAGAT AGTAG CC-3’), Reverse (5’-TCCAG CAGGA AGAAG ACAGG-3’); Nolz-1 null-Forward (5’-GTCGC CTTCC TCAGA AGCTA-3’), Reverse (5’-GATCT GGACA GGGAA AAGCA-3’); LacZ Forward (5’-CGTGG CCTGA TT CAT TCC-3’), Reverse (5’-ATCCT CTGCA TGGTC AGGTC -3’).

### Dissection of lung tissue and lung explant culture

Time-pregnant mice or adult mice were anesthetized with isoflurane. Lung tissues were dissected from E11.5, E12.5, E13.5, E14.5, E16.5, E18.5, P0, P7, P14, and adult mice. The esophagus and surrounding organs were removed from dissected lung tissue. The dissected E11.5-E13.5 lung tissues were placed on top of the culture insert (0.4 μm pore size, EMD Millipore). Explants were cultured with the BGjb medium (Invitrogen) supplemented with ascorbic acid (0.1 mg/ml) for 48 hr. For the rescue experiments, lung tissues were dissected from E11.5-E12.5 wild type or *Nolz-1* null mutant embryos. The lung explants were cultured with recombinant human Wnt2 protein (150 ng/ml, Novus Biologicals). For the induction of Nolz-1 experiments, the lung tissues were dissected from E13.5 wild type embryos. The lung explants were cultivated with recombinant mouse Fgf9 protein (200 ng/ml, R & D Systems, Inc.) or Fgf10 protein (200 ng/ml, R & D Systems, Inc.). Lung explants were cultured for 48 h with the change of fresh medium containing the recombinant proteins 24 h after cultivation. For each experimental condition, at least 4 lung explants from different embryos were used.

### Mesenchymal cell culture of lungs

The culture of lung mesenchymal cells was performed according to Lebeche et al. (Lebeche et al., 1999). The lungs of E14.5 littermate embryos were dissected, and were then digested with collagenase solution containing 0.1% collagenase (Sigma) and 0.01% DNAase (Sigma) in PBS at 37°C for 30 min. Enzymatic digestion was stopped by adding 2% fetal bovine serum (FBS). The digested lung tissue was triturated by pipetting 30 times with pipet-aid. The dissociated cells were spun at 1000 rpm for 5 min at room temperature, and the cell pellet was resuspended in 10 ml of DMEM supplemented with 2% FBS. The cells were plated with the density of 2 x 106 cells in 6-well dishes, and were then incubated with 5% CO_2_ at 37°C for differential cell adhesion. After 50 min of incubation, the medium containing epithelial cells was removed by washing twice with PBS followed by adding fresh medium while the mesenchymal cells remained adherent to the culture plates. The enrichment of mesenchymal cells in the culture was validated by immunostaining of Nolz-1 and TTF-1. The immunostaining indicated that less than 1% of the cultured cells were TTF-1-positive epithelial cells and 97% of the cultured cells were Nolz-1-positive mesenchymal cells (Figure S5).

### Electroporation of cell culture

E14.5 lung tissue was dissociated into single cells (5 x 106). Cells were re-suspended in 50 μl of electroporation solution containing 6 μg DNA plasmids, 2 mM CaCl2 and 55 mM glucose in PBS and were then transferred to an electroporation cuvette (2 mm, BTX). Electroporation was performed with an electroporator (170 V, 5 ms of duration, 1 pulse, BTX ECM 830). Electroporated cells were plated at the density of 4 x 106 cells in 6-well plates. The cells were cultured to enrich mesenchymal cells as described above. The cells were harvested 48 hr after seeding for qRT-PCR analysis and chromatin immunoprecipitation (ChIP) assay.

### Immunohistochemistry

E11.5, E12.5, E13.5, E14.5 whole embryos and E16.5, 18.5 whole lungs were fixed with 4% paraformaldehyde (PFA) overnight and were embedded with paraffin before sectioning at 5 μm. For frozen sections, after cryoprotection with 30% sucrose for two days, embryos were sectioned with a cryostat (Leica) at the thickness of 10 μm. For histological analysis, sections were stained with hematoxylin and eosin (H&E). For immunohistochemistry, paraffin-embedded sections were dewaxed by heating to 65°C for 30 min and then immersed in xylene for 3 x 10 min. The dewaxed sections were rehydrated by successive immersion in 100% ethanol x 5 min, 80% ethanol x 5 min, and 70% ethanol x 5 min followed by rinses in PBS. For cryosections, antigen retrieval was performed as needed. Briefly, sections were incubated with retrieval buffer (10 mM citrate acid, pH 6.0/0.05% Tween 20) at 95°C for 10 min. For immunohistochemistry, the sections were pretreated with 0.2% Triton X-100 in 0.1M PBS and 10% methanol/3% H2O2. After washing with PBS, sections were incubated in a blocking solution containing 3% serum in 0.1M PBS for 1 hr. The sections were incubated with primary antibodies diluted in 0.1 M PBS containing 1% serum, 0.2% Triton X-100 and 0.1% sodium azide overnight at room temperature or for two days at 4°C. The following primary antibodies were used: rabbit anti-Nolz-1 (1:1000) (Ko et al., 2013), mouse anti-TTF-1 (1:500, Cell Signaling), rabbit anti-Sox9 (1:2000, Abcam), rabbit anti-Axin2 (1:500, Abcam), mouse anti-β-catenin (1:1000, BD Biosciences), rat anti-BrdU (1:1000, EMD Millipore), mouse anti-Ki67 (1:500, Novocastra/Leica), rabbit anti-PH3 (1:1000, EMD Millipore), rabbit anti-active Caspase3 (AC3) (1:500, Cell Signaling), mouse anti-cyclin D1 (1:1000, Santa Cruz), rabbit anti-SP-C (1:500, Santa Cruz), mouse anti-CC-10 (1:500, Santa Cruz), rabbit anti-Aquaporin 5 (Aqp5) (1:500, Abcam), mouse anti-T1-alpha (1:500, Abcam), mouse anti-SMA (1:2000, Sigma), goat anti-SM22α (1:500, Santa Cruz), rat anti-PECAM-1 (1:500, Santa Cruz) and mouse anti-myc (1:1000, EMD Millipore), rabbit anti-Foxp1 (1:2000, Abcam), rabbit anti-Foxp2 (1:2000, Abcam). Secondary antibodies conjugated with chromophore Alexa-488 or Alexa-555 were used. For immunostaining of Nolz-1, TTF-1, AC3, and Ki67 immunostaining signals were further amplified by tyramide-FITC or tyramide-Cy3 (1:2,000, Perkin-Elmer, Wellesley, MA) after avidin-biotin complex incubation (Elite ABC kit, Vector Laboratories). Immunostained sections were counterstained with DAPI (1 μg /ml, Sigma).

### Whole-mount LacZ staining

E14.5 *Nolz-1+/+;BATGAL*, *Nolz-1−/−;BATGAL*, *Nolz-1+/+;TOPGAL* and *Nolz-1−/−;TOPGAL* lungs were dissected in ice-cold PBS, briefly washed in PBS twice, and fixed with 4% paraformaldehyde for 1 hr. The tissues were then pre-incubated with LacZ staining solution (150 mM NaCl, 1 mM MgCl2, 3.5 mM potassium ferrocyanide, 3.5 mM potassium ferricyanide, 0.3 mM chloroquine, 0.01% sodium deoxycholate, 0.2% NP-40 in 0.1M PB) for 1 hr at 37°C followed by incubation in LacZ staining solution containing X-gal (1 mg/ml) for 16 hr at 37°C. After the whole-mount LacZ stained lungs were photographed, tissues were cryosectioned for further analysis.

### *In situ* hybridization with digoxigenin-labeled probes

E11.5 and E13.5 lung sections were mounted on the slides coated with gelatin and poly-D-lysine (Sigma). The pCMV-Sport6-Wnt2 plasmid (IMAGE Id: 4162686) was linearized with the restriction enzyme EcoRI to generate the antisense probe. The digoxigenin-labeled antisense cRNA probe was synthesized by in vitro transcription with T7 polymerase (Promega), RNA labeling mix (Roche), and the linearized pCMV-Sport6-Wnt2 plasmid. After fixation with 4% PFA and rinsing with PBS, the sections were treated with 0.2 N HCl and proteinase K (10 μg/ml), and were then prehybridized with 50% formamide in 2 X SSC for 90 min at 65°C. The sections were hybridized with the Wnt2 cRNA probe solution (10.6% dextran sulfate, 53% formamide, 1 mM EDTA, 10.6 mM Tris-HCl pH 8.0, 318 mM NaCl, 1.06 X Denhart’s solution, 200 μg/ml tRNA and 10 mM DTT) for 16 hr at 65°C. Sections were then washed with 2 X SSC containing 50% formamide for 1 hr at 65°C, and were then treated with RNase A (20 μg /ml). The sections were washed with 2 X SSC and 0.2 X SSC for 20 min followed by incubation in a blocking solution containing 2% blocking reagent (Roche) and 20% sheep serum for 1 hr. The sections were incubated with horseradish peroxidase (HRP) conjugated sheep anti-digoxigenin antibody (1:100, Roche) overnight at RT. The signals were detected using a TSA amplification kit (PerkinElmer) following the manufacturer’s protocol. Sections were counterstained with DAPI.

### *In situ* hybridization with isotope-labeled probes

The pGEM-T-Nolz-1 plasmid was linearized with SalI to generate antisense probes. The production of cRNA probes was carried out in the presence of 35S-UTP (NEN) with T7 RNA polymerase by in vitro transcription. Brain sections were incubated with 10 μg/ml proteinase K at 37°C for 5 min, RNA hybridization was performed with the 35S-UTP-cRNA probe (107 cpm/ml) in the hybridization buffer containing 50% formamide, 10% dextran sulfate, 0.3 M NaCl, 1x Denhart’s solution, 0.01 M Tris (pH8.0), 1 mM EDTA. 0.5 mg/ml yeast tRNA, and 1 mM DTT at 58°C for 16 hr. After several rinses in 4x SSC, the slides were treated with RNase A (10 μg/ml) and washed with 2x SSC, 1x SSC, and 0.5x SSC at room temperature. The slides were then washed with 0.1x SSC at 50 for 30 min followed by another 0.1X SSC wash at room temperature for 5 min. Ethanol dehydration was carried out with serial rinses with 50%, 75%, 95% ethanol, and then 100% ethanol twice, each for 3 min. The slides were vacuum-dried for 1 hr, and were exposed to X-ray film. Signals were detected by autoradiography.

### BrdU incorporation assay

Time-pregnant mice were injected with 5-bromo-2’deoxyuridine (BrdU) (50 mg/kg body weight, Sigma) 1 h before the harvest of embryos. Embryos were sectioned and processed for immunohistochemistry as described above. The lung explant culture was incorporated with BrdU (10 µM) for 1 hr and then harvested for further analysis.

### TUNEL assay

The TUNEL assay was performed in cryosectioned lung tissue using the *In Situ* Cell Death Detection Kit (Roche) according to the manufacturer’s instructions. Briefly, sections were treated with 0.1 M glycine in PBS buffer for 30 min and then incubated with 0.1% Triton X-100 in 0.1% sodium citrate buffer for 3 min at 4°C. Following 3 x 5 min rinses with PBS, the sections were incubated with the solution containing 10% terminal transferase (TdT) for 1 hr at 37°C to label the free 3’-OH ends in fragmented DNA with fluorescein-conjugated dUTP. The sections were counterstained with DAPI.

### Quantitative RT-PCR

Lung tissues and cultured cells were lysed in Trizol reagent (Invitrogen), followed by RNA extraction. A two-step quantitative reverse transcription-polymerase chain reaction (qRT-PCR) was performed. Total RNA (2 μg) was mixed with oligo-dT (Roche) and dNTP (Yeastern, Taiwan) and then denatured at 65°C for 5 min. After the addition of 0.1 M DTT, RNase inhibitor (RNasin, Promega), 5X buffer, and SuperScrip III (Invitrogen) transcriptase, reverse transcription of RNA into cDNA were carried out at 50 ° C for 2 hr. The reaction was stopped by incubation at 70°C for 15 min. The cDNA was used for the real-time PCR reaction (Applied Biosystems StepOne system). The PCR reaction solution contained ∼1 ng of cDNA and 2X TaqMan® Fast Universal PCR Master Mix (Applied Biosystems®). PCR reactions were carried out under the following conditions: 95 ° C for 10 min followed by 40 cycles at 95°C for 10 sec, 60°C for 1 min with the following sets of PCR primers: Aquaporin-5-Forward (5’-ATGAA CCCAG CCCGA TCTTT-3’), Reverse (5’-ACGAT CGGTC CTACC CAGAA G-3’); Axin2-Forward (5’-CTCCC CACCT TGAAT GAAGA-3’), Reverse (5’-ACATA GCCGG AACCT ACGTG-3’); Bmp4-Forward (5’-GCCAT TGTGC AGACC CTAGT-3’), Reverse (5’-ACCCC TCTAC CACCA TCTCC-3’); c-myc-Forward (5’-TCGTG AGAGT AAGGA GAA-3’), Reverse (5’-CAAGG TTGTG AGGTT AGG-3’); cyclin D1-Forward (5’-TGTTC GTGGC CTCTA AGATG AAG-3’), Reverse (5’-AGGTT CCACT TGAGC TTGTT CAC-3’); Fgf9-Forward (5’-TGCAG GACTG GATTT CATTT AGAG-3’), Reverse (5’-CGAAG CGGCT GTGGT CTTT-3’); Fgf10-Forward (5’-GAGAA GAACG GCAAG GTCAG-3’), Reverse (5’-CCCCT TCTTG TTCAT GGCTA-3’); Fgfr2-Forward (5’-AAGAT GATGC CACAG AGA-3’), Reverse (5’-CCAGG AGGTT GATAA TGTTC-3’); Foxj1-Forward (5’-TGGAC TGGCT CCATT TTA-3’), Reverse (5’-ATGCA TTTAG ACACC GGA-3’); GAPDH-Forward (5’-CGTGG AGTCT ACTGG TGTCT TC-3’), Reverse (5’-TGCAT TGCTG ACAAT CTTGA G-3’); Gli2-Forward (5’-TCCCT CAGCA AATGG AAGTT G-3’), Reverse (5’-CCGTG CTCCC GTTGA TG-3’); Gli3-Forward (5’-GTGTG ACTTC TCCTC TTA-3’), Reverse (5’-GCATC CTTCC TATTA CCT-3’); Lef1-Forward (5’-TTACT CTGGC TACAT AATGA T-3’), Reverse (5’-ACGGG CACTT TATTT GAT-3’); Mycn-Forward (5’-GGAAG TTCAC ACCTA AGT-3’), Reverse (5’-AGTTA TGTAT CAGCG TCAT-3’); Nolz-1-Forward (5’-TTACT CTGGC TACAT AATGA T-3’), Reverse (5’-GGCTC CTTCT TATCT GAACC-3’); Pdgfrα-Forward (5’-TGGCA TGATG GTCGA TTCTA-3’), Reverse (5’-CGCTG AGGTG GTAGA AGGAG-3’); Pdgfrβ-Forward (5’-CCGGA ACAAA CACAC CTTCT-3’), Reverse (5’-TATCC ATGTA GCCAC CGTCA-3’); Pdgfc-Forward (5’-CTTAG TTGTC TTGAT ATGG-3’), Reverse (5’-GAGCA TCTGT CTATC TAT-3’); Scgb1a1-Forward (5’-TCCTA ACAAG TCCTC TGTGT AAGA-3’), Reverse (5’-AGGAG ACACA GGGCA GTGAC A-3’); Shh-Forward (5’-CCCAA TTACAA CCCCG ACAT-3’), Reverse (5’-GTCTT TGCAC CTCTG AGTCA TCA-3’); SMA-Forward (5’-TGGCA TCAAT CACTT CAA-3’), Reverse (5’-CCTAT CTGGT CACCT GTA-3’); SM22α-Forward (5’-GCCCA GCCTC TACAT CTT-3’), Reverse (5’-GAATG CTAAC AGGAG TGACA A-3’); SftpA-Forward (5’-CTCCA GACCT GTGCC CATAT G-3’), Reverse (5’-ACCTC CAGTC ATGGC ACAGT AA-3’); SftpB-Forward (5’-ACGTC CTCTG GAAGC CTTCA-3’), Reverse (5’-TGTCT TCTTG GAGCC ACAAC AG-3’); SftpC-Forward (5’-ACCCT GTGTG GAGAG CTACC A-3’), Reverse (5’-TTTGC GGAGG GTCTT TCCT-3’); T1-α-Forward (5’-AGGTA CAGGA GACGG CATGG T-3’), Reverse (5’-CCAGA GGTGC CTTGC CAGTA-3’); Tgf-β2-Forward (5’-TATTG ATGGC ACCT CTAC-3’), Reverse (5’-ACAAC ATTAG CAGGA GAT-3’); Wnt2-Forward (5’-TCTTG AAACA AGAAT GCAAG TGTCA-3’), Reverse (5’-GAGAT AGTCG CCTGT TTTCC TGAA-3’); Wnt2b-Forward (5’-GCAGA CAACA GACTA GATTC-3’), Reverse (5’-TACGA TTGGA TGAAG AGGAA-3’); Wnt7b-Forward (5’-GCAGT GTGGA TGGAT GTT-3’), Reverse (5’-GGCTA CCCAG TCGTT AGTA-3’).

### Western blot

The lung tissues from E12.5, E14.5, E16.5, E18.5 embryos, and P0, P7, P14 and adult mice were lysed in RIPA buffer [20 mM HEPES, pH 7.8, 150 mM NaCl, 1 mM EDTA, 1% NP40, protease inhibitor cocktail (Roche Applied Science]. Tissue lysates (20 μg) of embryonic lungs were subjected to SDS-PAGE and transferred to a polyvinylidene difluoride membrane (PVDF; Millipore). Immunoblotting was performed using an anti-Nolz-1 antibody (1:1000) and anti-actin (1:10000, Sigma). Secondary antibodies were HRP-conjugated goat anti-rabbit IgG and goat anti-mouse IgG (Jackson ImmunoResearch Laboratories). For testing the specificity of the Nolz-1 antibody, the anti-Nolz-1 antibody was incubated with 50 μg Nolz-1 recombinant protein overnight at 4°C. The preabsorbed antibodies were then used for immunoblotting. Immunoblotting ECL signals were detected with a Luminescence Imaging System (Fuji Film LAS-4000).

### Chromatin immunoprecipitation (ChIP) assay

The pcBIG-myc-Nolz-1-ires-EGFP plasmids were electroporated into lung mesenchymal cell culture as described above. The ChIP assay was performed according to the manufacturer’s instructions (EZ-Magna ChIPTM A/G kit, EMD Millipore). In brief, 1/10 volume of fresh 11% formaldehyde solution was added to culture plates for 10 min at room temperature followed by quenching formaldehyde with 1/20 volume of 2.5 M glycine. After washing twice with 5 ml of PBS, cells were harvested by spinning at 1,350 x RPM for 5 min at 4°C. Cells were re-suspended in 0.5 ml of cell lysis buffer containing 2.5 μl of protease inhibitor cocktail II. Cells were spun down at 800 g for 5 min at 4°C. The pellet was resuspended in 0.5 ml nuclei lysis buffer and subjected to sonication to shear DNA with a sonicator (5s on, 30 s off, 4 cycles, power 4, 2 rounds, MISONIX Inc.). The debris was removed by centrifuge at 13,000 rpm for 10 min at 4°C. Mouse anti-myc antibody (1 μg) or mouse IgG (1 μg) (EMD Millipore) were incubated with protein A/G Magna beads (20 μl) and the sheared DNA lysate overnight at 4°C. Antibody-DNA complexes were washed sequentially with low salt, high salt, LiCl and TE buffers followed by incubation in the ChIP elution solution containing proteinase K at 65°C for 2 hr. DNA purification was performed using spin columns. The purified DNA products were used as templates for PCR reaction with N1 primers: Forward (5’-GAGGA GCTGT GTGTG CGTAA-3’), Reverse (5’-CCAGT GTCTG ACCAG GTTGA-3’) and N2 primer: Forward (5’-CAGGT CAGAA GGGGT GTTTG-3’), Reverse (5’-GCCTC CTAGA AAAGG GATGG-3’). PCR was performed with the following reaction condition: 95°C for 10 min followed by 30 cycles at 95°C for 10 sec, 58°C for 10 sec and 72°C for 20 sec. An additional 72°C was run for 3 min at the end of the PCR reaction. The PCR amplicons of the ChIP assay were analyzed by 2% agarose gel electrophoresis.

### Constructions of plasmids

The genomic region of the mouse Wnt2 gene (NM_023653.5) containing the putative Nolz-1 binding motif N1 (-722 to -904, +1 translation start site) and the N2 motif (-2149 to - 2335) was amplified by PCR with the N1-ChIP and N2-ChIP primers. The amplicons of N1 (183 bp) and N2 (187 bp) were constructed into the pGL3-basic plasmid (Promega).

### Luciferase reporter and TOPFlash/FOPFlash assays

Primary lung mesenchymal cells were transfected with pGL3-N1 (or pGL3-N2), pcBIG-myc-Nolz-1-ires-EGFP (or control pcBIG-ires-EGFP) and pGL4.74-Renilla luciferase plasmids in a ratio of 1:1:0.1 (4 μg/6-well dishes) by using Lipofectamine LTX PLUSTM (Invitrogen). The luciferase activity was assayed using the Dual-Luciferase Reporter Assay Kit (Promega). The luciferase signals were detected with a luminescence reader (VICTORTM x2, PerkinElmer). The TOPFlash or FOPFlash plasmids were cotransfected with pcBIG-myc-Nolz-1-ires-EGFP (or mock control pcBIG-ires-EGFP) and pGL4.74-Renilla luciferase plasmids in a ratio of 0.5:1:0.1 (4 μg/6-well dishes) using Lipofectamine LTX PLUS (Invitrogen). The luciferase activity was assayed using the Dual-Luciferase Reporter Assay Kit (Promega). The luciferase signals were detected with a luminescence reader (VICTORTM x2, PerkinElmer).

### Quantitative and statistical analysis

Photomicrographs of the dissected lungs were taken under a dissecting microscope (Nikon SMZ800) with the aid of a digital camera (Nikon DXM 1200C). Photomicrographs of lung histology were acquired with the aid of fluorescence microscopes (BX51, BX63, Olympus) and a confocal microscope (Zeiss LSM700). Photomicrographs of lung sections and lung explant cultures were selected for morphological quantification. At least 3 pairs of wild type and *Nolz-1* null mutant lungs from independent littermates were used for analysis. The number of epithelial branches of the left lobe in lung explants at 48 hr after culture was normalized with the number of epithelial branches at 0 hr. The numbers of Sox9+, BrdU+, PH3+ and cyclin D1+ cells were counted using ImageJ software (NIH, USA). The intensity of PCR amplicons of the ChIP assay was measured by ImageJ software. All quantitative data were first assessed for normality. For the normally distributed data, Student’s two-tailed t test was used for all statistical analyses. Error bars represent Mean ± SEM for at least three independent experiments.

## Acknowledgments

We thank Drs. C.-M. Chen and L.-R. You for consultation of embryonic lung dissection, providing reagents, transgenic mice and insightful discussion, and the NYMU Genomic Research Center for providing cDNA clones. This work was supported by the Ministry of Science and Technology grants NSC99-2311-B-010-005-MY3, NSC102-2321-B-010-018, MOST103-2321-B-010-009, the Ministry of Science and Technology-Taiwan grants MOST107-2320-B-010-041-MY3, MOST110-2320-B-A49A-532-MY3, MOST110-2326-B-A49A-504 (F.-C.L.), Postdoctoral Fellowship grant MOST109-2811-B-010-538, MOST110-2811-B-A49A-031 (S.-Y. C) and the Higher Education Sprout Project by the Ministry of Education in Taiwan (F.-C.L.).

## Author Contributions

S.-Y.C. and F.-C.L. designed experiments; S.-Y.C. performed experiments; S.-Y.C. and F.-C.L. analyzed the data and wrote the manuscript; F.-C.L. supervised the project.

