## Supplemental information for "Fgf9-Nolz-1-Wnt2 Signaling Axis Regulates Morphogenesis of the Lung"

**This Supplemental Information includes:**

Supplemental Figures: Figures S1 to S5

Figure S1, Related to Figures 2, 3, 4, 5, 6, 7
Figure S2, Related to Figure 2
Figure S3, Related to Figure 3
Figure S4, Related to Figure 2, 4
Figure S5, Related to Figure 5

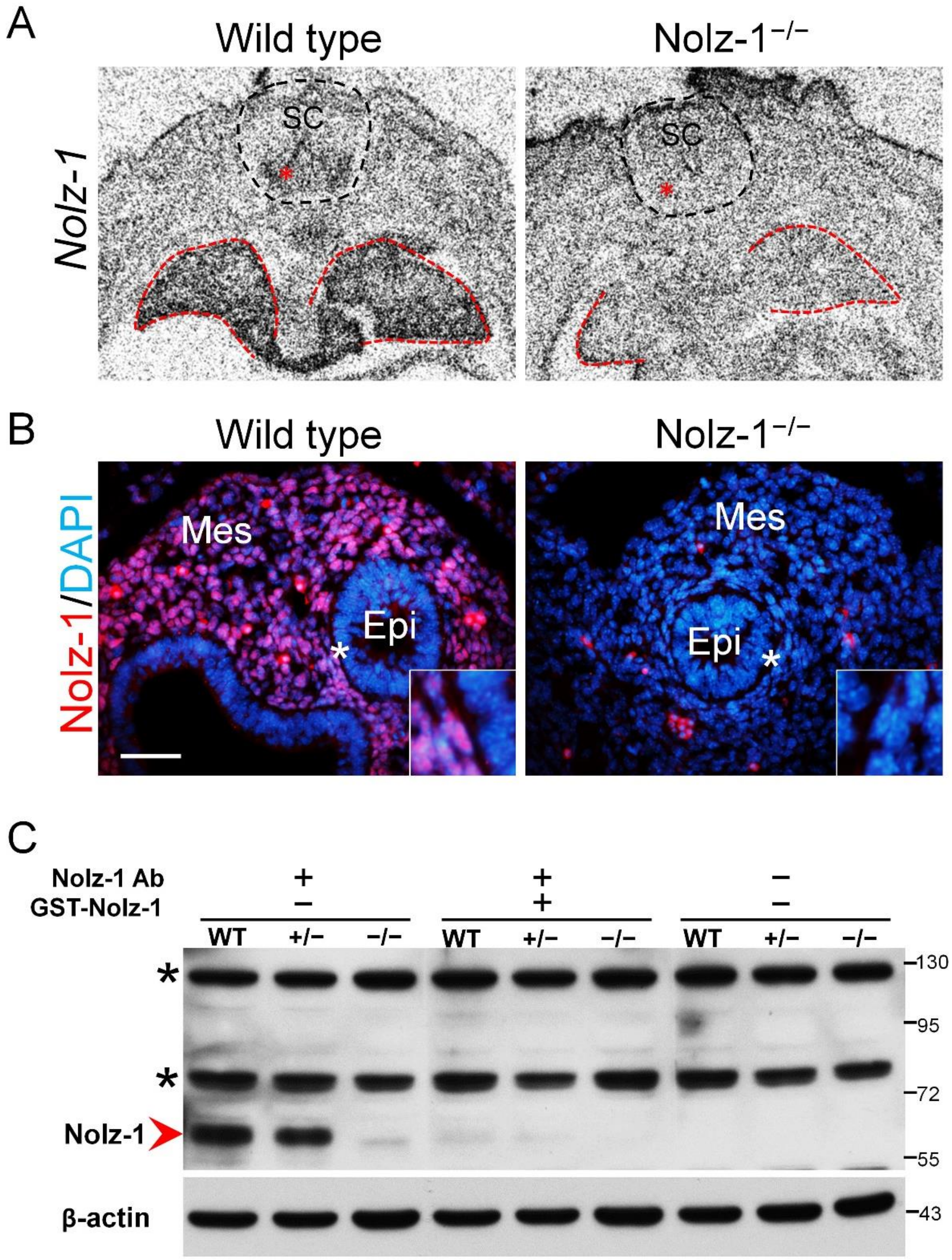


**Figure S1. Confirmation of loss of *Nolz-1* mRNA and protein in *Nolz-1* null mutant lungs.** (A) *In situ* hybridization confirms that *Nolz-1* mRNA is absent in E12.5 *Nolz-1* mutant lung (red dotted line) and spinal cord (red asterisks, motor neuron). SC: spinal cord. (B) Immunocytochemistry shows that Nolz-1 protein is not expressed in the E12.5 *Nolz-1* mutant lung. The insets show high magnification of the regions indicated by asterisks. Epi: epithelium; Mes: mesenchyme. (Scale bar, 50 μm). (C) Immunoblotting shows the loss of Nolz-1 protein (arrowhead in red) in *Nolz-1* mutant lung. The asterisks indicate non-specific bands.

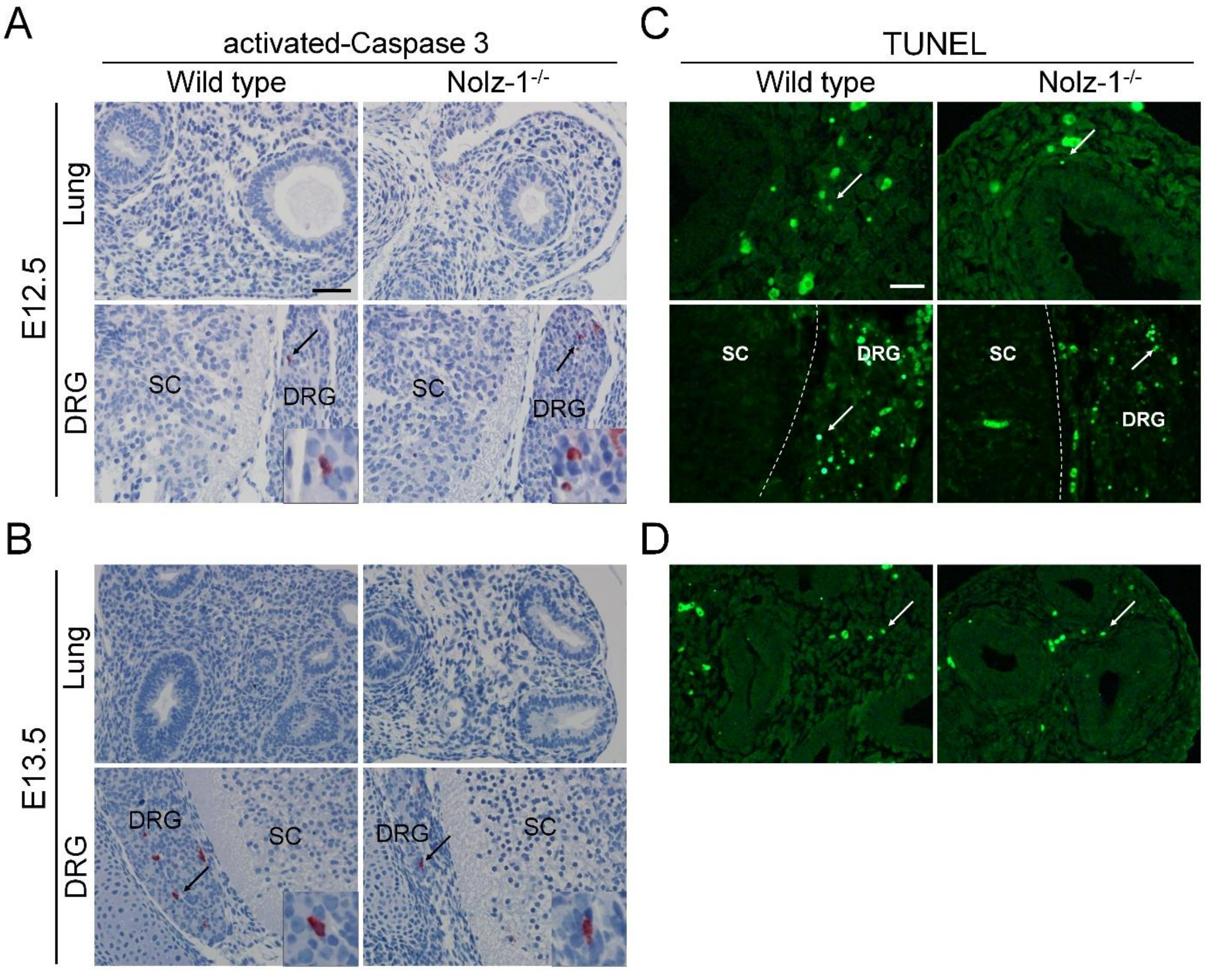


**Figure S2. Absence of abnormal apoptosis in early stages of *Nolz-1* null mutant lungs.** (A and B) Immunostaining of active Caspase-3 (AC3) shows no significant difference between wild type and Nolz-1 mutant lungs at E12.5 (A) and E13.5 (B). The AC3 immunostaining is validated by positive controls in which many AC3-positive cells are present in the dorsal root ganglion (DRG) of the same sections at E12.5 and E13.5. (Scale bars, 50 μm). (C and D) Similar results are obtained using the terminal deoxynucleotidyl transferase-mediated dUTP nick end labeling (TUNEL) assay. Few TUNEL-positive cells are present in the lungs of wild type and *Nolz-1* mutant without a significant difference at E12.5 (C) and E13.5 (D). Some TUNEL-positive cells are present in the DRG of the same sections of E12.5 embryos as a positive control. (Scale bars, 20 μm).

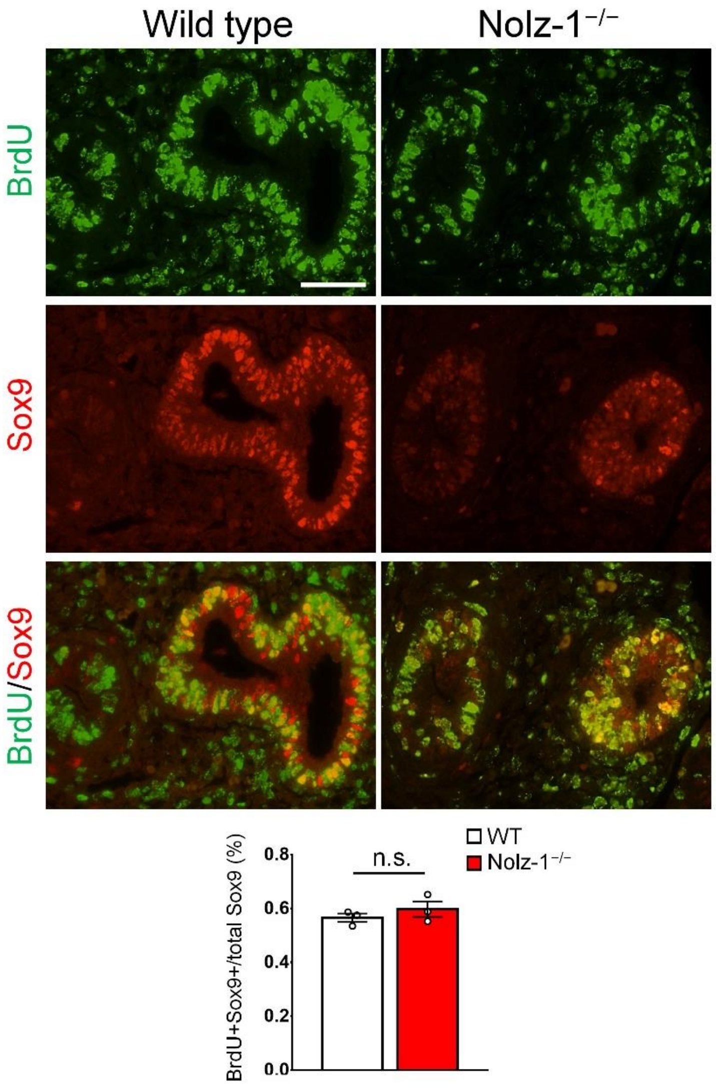


**Figure S3. The normal proliferation of Sox9-positive distal epithelial cells in *Nolz-1* null mutant lungs.** Double immunostaining of BrdU and Sox9 show similar results between wild type and mutant lungs. Quantification of the density of $BrdU^+Sox9^+$ double-positive cells in the epithelium, there is no significant difference between wild type and mutant lungs. Student's *t*-test, n.s. not significant, n = 3. (Scale bars, 50 μm).

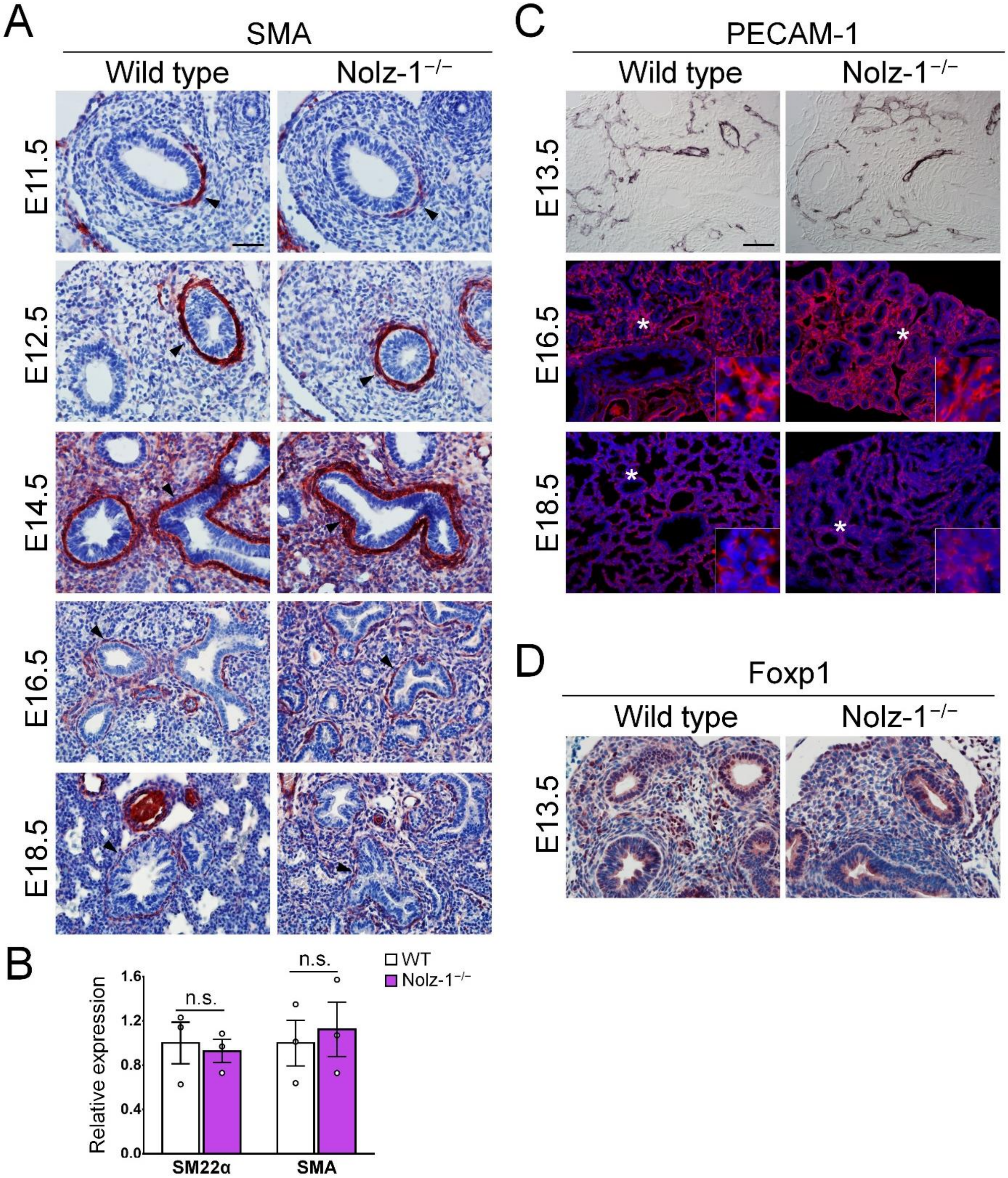


**Figure S4. The effects of *Nolz-1* null mutation on differentiation of mesenchymal cells in developing mouse lungs.** (A) Immunostaining shows a decrease of smooth muscle actin (SMA) in *Nolz-1* mutant lung at E11.5, but is not changed at E12. 5, E14.5, E16.5 and E18.5 between wild type and mutant lungs. Arrowheads indicate smooth muscle layers in E11.5-E18.5 lungs. (Scale bars, 50 μm). (B) The qRT-PCR assay shows

*SMA* and *SM22α* mRNA, progenitor marker of smooth muscle cells, is not changed in the E12.5 *Nolz-1* mutant lung. Student's *t*-test, n.s. not significant, n = 3. (C) The immunostaining pattern of PECAM-1/CD31, a marker of vascular endothelial cells appears similar between wild type and *Nolz-1* mutant lungs at E13.5, E16.5 and E18.5. The insets show high magnification of the regions indicated by asterisks. (Scale bars, 100 μm). (D) The expression pattern of Foxp1, which is expressed in the mesenchyme and distal parts of epithelium, is not changed in E13.5 mutant lungs. (Scale bars, 100 μm).

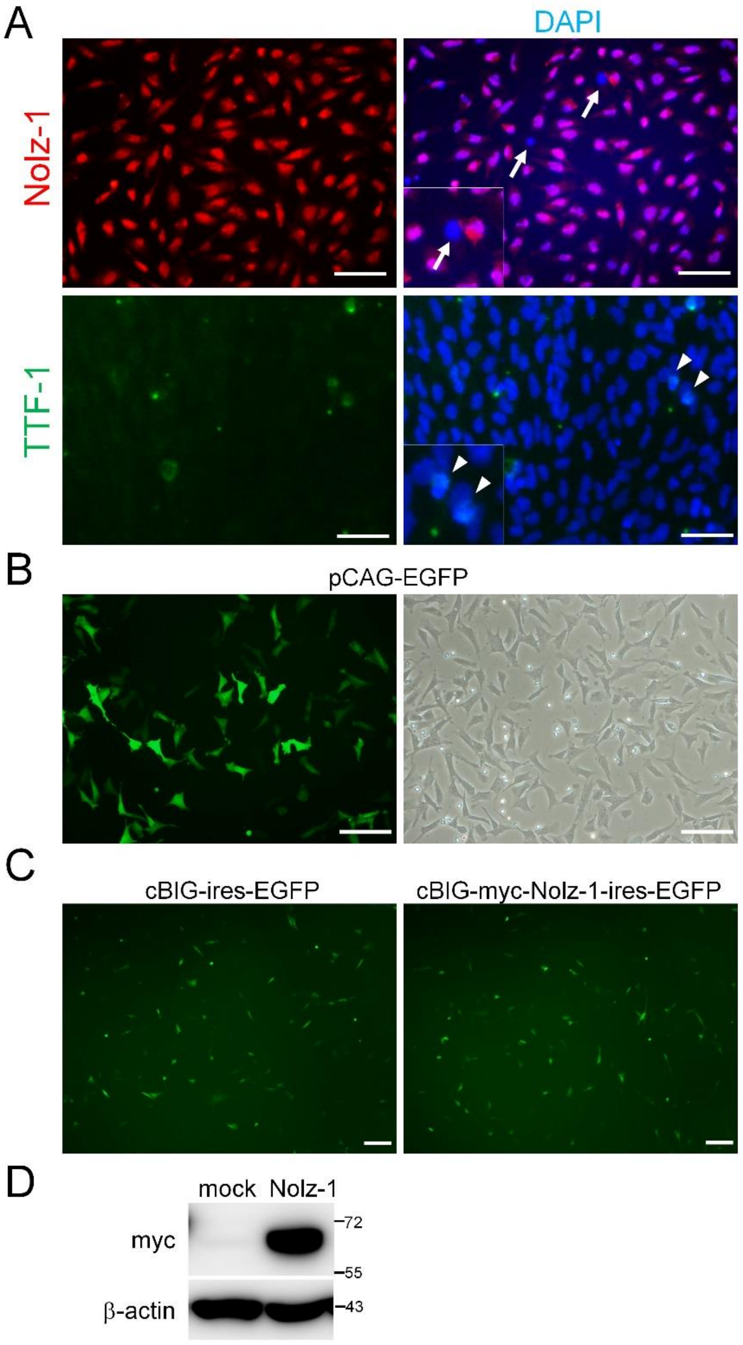


**Figure S5. Characterization of primary mesenchymal cells of developing lungs.** (A) The cultured lung mesenchymal cells are validated by immunostaining of Nolz-1 and TTF-1. The Nolz-1-negative cells are indicated by arrows. The immunostaining indicates that less than 1% of the cultured cells are TTF-1-positive epithelial cells (arrowhead) and 97 % of the cultured cells are Nolz-1-positive mesenchymal cells. (Scale bar, 50 μm). (B) Evaluate the efficiency of gene delivery by electroporation of pCAG-EGFP plasmid. About

40% of cells express GFP after electroporation. (Scale bar, 50 μm). (C) Electroporation of cBIG-ires-EGFP (mock control) and cBIG-myc-Nolz-1-ires-EGFP plasmids into primary lung mesenchymal cells. (Scale bar, 50 μm). (D) Overexpression of myc-tagged Nolz-1 protein in primary lung mesenchymal cells is demonstrated by Western blotting.
